# Specific light-regime adaptations, terpenoid profiles and engineering potential in ecologically diverse *Phaeodactylum tricornutum* strains

**DOI:** 10.1101/2024.08.05.606631

**Authors:** Luca Morelli, Payal Patwari, Florian Pruckner, Maxime Bastide, Michele Fabris

**Affiliations:** SDU Biotechnology, Faculty of Engineering, University of Southern Denmark, Campusvej 55, Odense M, DK-5230, Denmark; SDU Climate Cluster, Faculty of Science, University of Southern Denmark, Campusvej 55, Odense M, DK-5230, Denmark

**Keywords:** *Phaeodactylum tricornutum*, light regimes, photosynthesis, terpenoids, genetic transformation

## Abstract

Microalgae, and among them, the diatom *Phaeodactylum tricornutum* stand out with their remarkable versatility and metabolic engineering potential. Diatoms exhibit substantial variability in metabolism, photosynthetic physiology and environmental adaptation, even across the same species. These factors can affect the design and outcome of metabolic engineering strategies. In this study, we profiled the diversity of biotechnologically relevant traits of three *P. tricornutum* strains (Pt1, Pt6, and Pt9) under different light regimes to identify the most suitable chassis to be employed as bio-factory to produce high-value terpenoids. We conducted detailed assessments of these strains, using pulse amplitude modulated (PAM) fluorometry to measure photosynthetic efficiency and we analyzed the composition of pigments and triterpenoids, as main terpenoid metabolic sinks. Parameters such as the maximum quantum yield of PSII (Fv/Fm), the efficiency of excitation energy capture (Fv’/Fm’), and OJIP kinetics were used to estimate photosynthetic performance in different light regimes. Additionally, we evaluated their transformation efficiency and their capacity to produce heterologous monoterpenoids, using geraniol as a model product. Our findings revealed that Pt1, widely used in laboratories, exhibits robust growth and photosynthetic performance under standard laboratory conditions. Pt6, adapted to intertidal environments, shows unique resilience in fluctuating conditions, while Pt9, with its high-temperature tolerance, excels under continuous high irradiance. Additionally, this variability across strains and light conditions influenced the metabolic output of each strain. We concluded that understanding the physiological responses of different *P. tricornutum* strains to light is crucial for optimizing their use in metabolic engineering. The insights gained from this research will facilitate the strategic selection and exploitation of these strains in algae biotechnology, enhancing the production of commercially valuable compounds such as high-value terpenoids and derivatives. This comprehensive characterization of strains under varying light conditions offers a pathway to more efficient and targeted metabolic engineering applications.

**Highlights:**

- Pt1, Pt6, and Pt9 exhibit distinct physiology under different light regimes.
- Pt9 is photosynthetically more performant in continuous light, Pt6 in photoperiod.
- Light regimes affect pigments and triterpenoid content in all three strains.
- Each strain exhibits a specific carotenoid and triterpenoid composition.
- Bacterial conjugation of episomes varies across strains, it is more efficient in Pt1.
- Pt1 is suited for the heterologous synthesis of monoterpenes (geraniol).

## 1. Introduction

In the search for sustainable microbial platforms to produce valuable compounds, microalgae have emerged as compelling candidates for their versatility, ubiquitous distribution, and suitability for genetic and metabolic engineering. Photosynthetic eukaryotic microalgae have colonized nearly all and the most diverse habitats worldwide, and showcase remarkable biological diversity with over 200,000 reported species [1]. In contrast to other microbes commonly used in bio-manufacturing, such as yeasts (e.g., *Saccharomyces cerevisiae* or *Pichia pastoris*) and bacteria (e.g. *Escherichia coli*), phototrophic microalgae possess the unique ability to utilize sunlight to fix carbon dioxide (CO2) through photosynthesis, reducing the dependence on fermentable sugars [2,3]. They can convert inorganic carbon into natural products, such as lipids, at a much faster rate than conventional oleaginous crops [4]. Moreover, microalgae often possess peculiar metabolic traits that sets them apart from conventional microbial hosts, and may offer new opportunities for industrial exploitation [5]. These metabolic processes, inherently reliant on photosynthesis, are profoundly influenced by factors like light intensity, regime, and quality. Thus, the modulation of light conditions emerges as a crucial determinant in providing optimal growth settings for microalgae.

Within microalgae, the pennate diatom *Phaeodactylum tricornutum* has garnered significant attention as a host for synthetic biology [6] boasting an extensive genetic toolkit encompassing differentiated and efficient transformation methods such as biolistic transformation, electroporation and conjugation, that can be used for random chromosomal integration of genetic constructs, or maintain them onto self-replicating, extrachromosomal episomes [7]. Additionally, it encompasses modular multi-gene DNA assembly [8] and genome editing technologies, rendering it an advanced platform for genetic manipulation [9].

Differently to hosts like *E. coli* and *S. cerevisiae*, and green algae such as *Dunaliella salina* and *Haematococcus pluvialis*—often cultivated on a large scale for their carotenoid accumulation— *P. tricornutum* possesses both the cytosolic mevalonate (MVA) and plastidial methylerythritol phosphate (MEP) pathways which synthesize precursors for multiple classes of terpenoid-based metabolites, including carotenoids, sterols, and tocopherols [10]

However, despite their promising attributes, the photosynthetic physiology of diatoms varies significantly among different strains of the same species, leading to a gradient of metabolic performances originating from diverse environmental adaptations [10,11]. For instance, diatoms possess fucoxanthin (Fx) as their primary light-harvesting xanthophyll. This compound is commercially valuable for its antioxidant properties [13]. In *P. tricornutum*, Fx is steadily accumulated during cultivation under low irradiances to maximize photon capture, and its levels are closely linked to the photosynthetic performance of the algae [14,15]. Furhermore, accumulation of Fx is also regulated by the expression of the *VDE1* gene, which shows differential expression across various strains of *P. tricornutum* (Li et al., 2024). Understanding the photo-physiological variability of this diatom is crucial in selecting the most suitable strain for metabolic engineering. One strain may be more compatible with artificial cultivation conditions than another, and a particular strain with a unique photosynthetic metabolism could exhibit elevated levels of specific metabolites, making it the best candidate for increasing the production of that metabolite of interest.

For its particular significance in ecology, research and industry, ten strains of *P. tricornutum*, catalogued as Pt1-Pt10 and sourced from distinct collections and geographic locations, were systematically characterized morphologically and genetically [15,16]. While this comprehensive effort elucidated the morphotypes and growth capabilities of multiple *Phaeodactylum* strains, a concurrent photosynthetic characterization is still lacking. Many studies have focused on the biomass and metabolite productivity of this diatom, particularly fatty acid and fucoxanthin accumulation under varying light and cultural conditions. However, these investigations often neglect the simultaneous assessment of photosynthetic performance and do not explore whether different strains show variations in transformation efficiency or recombinant protein production [13,17].

This study aims to address the gap by assessing the performance of three selected strains (Pt1, Pt6, and Pt9) identified by De Martino et al. These assessments were conducted under varying light regimes, reflecting conditions common in molecular biology research for diatom maintenance and culturing. Pt1, designated as accession CCMP632 in March 2001 and later identified as CCAP 1055/1, is a pivotal strain as one of its monoclonal cultures (i.e., Pt1 8.6) was chosen for the first genome sequencing project (http://genome.jgi-psf.org/Phatr2/Phatr2.home.html) [19], and it is the most widely used strain in routine laboratory experiments.

Pt6, deposited at CCMP in December 1986 with the accession number CCMP631, exhibits more frequently the oval morphotype [16], and it thrives in intertidal rock pools and water tanks, deviating from the typical planktonic habitat [20].

Finally, Pt9, collected in Micronesia in February 1981 and stored at CCMP under accession number CCMP633, displays unique characteristics. Unlike other strains, Pt9 thrives at higher temperatures (20°C to 26°C) and under substantially elevated irradiances. It also demonstrates morphological adaptability, transitioning from the typical fusiform shape to an oval shape at lower temperatures [16].

By employing pulse amplitude modulated (PAM) fluorometry, we systematically investigated chlorophyll as a variable fluorescence—an essential marker of photosynthetic efficiency. This technique involves subjecting diatom cultures to a modulated light source, inducing variations in chlorophyll fluorescence, which are captured and analyzed in real-time [20,21]. Parameters derived from this technique, including the maximum quantum yield of PSII (Fv/Fm), the efficiency of excitation energy capture by open PSII reaction centers (ϕPSII), and the OJIP kinetics representing sequential electron flow through PSII, serve as pivotal indicators reflecting the photosynthetic health of diatom strains. Furthermore, we investigated how different photoperiod affects the pigments and triterpenoid composition of various strains. Chlorophylls and carotenoids, crucial in light response, in fact, are also important in industry and pharmaceuticals as colorants and antioxidants [23]. Triterpenoids, including phytosterols, regulate cell membrane dynamics are valued as nutraceuticals for their cholesterol-lowering and anti-inflammatory properties [24]. The biosynthesis and metabolic fluxes of the precursors to carotenoids and triterpenoids are the starting point for every metabolic engineering strategy aimed at optimizing the production of these endogenous [25] and enable the biosynthesis of heterologous ones [26–28]. Yet, the regulation and activity of the terpenoid metabolism of diatom is elusive and analyzing the composition of the main terpenoid compounds in different strains will offer invaluable insights into their adaptability and potential suitability for applications in metabolic engineering. In this perspective, we also evaluated the transformation efficiency of each strain and their performance in producing heterologous geraniol, a commercially relevant plant-derived monoterpenoid that is a component of rose essential oil and used in the cosmetic industry, and precursor of high-value pharmaceuticals such as the monoterpenoid indole alkaloids vinblastine and vincristine [26].

The knowledge gained from this investigation holds significant potential in shaping the strategic selection and utilization of *P. tricornutum* strains for targeted metabolic engineering purposes. Ultimately, this study aims to facilitate an informed and optimized exploitation of these diatom strains within algae biotechnology.

## 2 Materials and methods

### 2.1 Algae cultures

Axenic cultures of *P. tricornutum* CCAP1055/1, CCAP1055/4 and CCAP1055/5 (respectively named Pt1, Pt6 and Pt9 as in De Martino et al., 2007) were obtained from the CCAP (Culture Collection of Algae and Protozoa, SAMS Research Services Ltd., Scottish Marine Institute, Oban, U.K., http://www.ccap.ac.uk) and were grown in liquid Enriched Artificial Sea Water (ESAW) medium [29]. For the experiment, diatoms were cultured in individual 50 ml flasks in fully controlled Innova ® S44i shaking incubators (Eppendorf) under 100 μE m^-2^sec^-1^ light at 21°C shaking at 95 rpm following different light regimes (*see “Light regimes treatment” section*). Algal stock cultures were maintained in a static incubator (MediLow-M) under the aforementioned conditions, with regular subculturing until the beginning of the experiment. Cell density was regularly assessed through a Guava H5 (Cytek) flow cytometer model while chlorophyll levels were assessed after proper extraction (*See “Pigment analysis” section*) using a Shimadzu UV-1280 spectrophotometer.

### 2.2 Light regime treatment

To assess the impact of light regimes on the photosynthetic performance, stock cultures were diluted to achieve an initial OD750 of 0.03 for all strains. Precultures of the three strains were cultivated under the specified conditions, following a light regimen of continuous light (CL) or one of 12 hours of light and 12 hours of darkness (PP). Parameters such as cell and chlorophyll amount and photosynthetic performance (see 2.3) were monitored at 3, 5, and 7 days from the start of the experiment (Fig. 1a).

**Figure 1:**
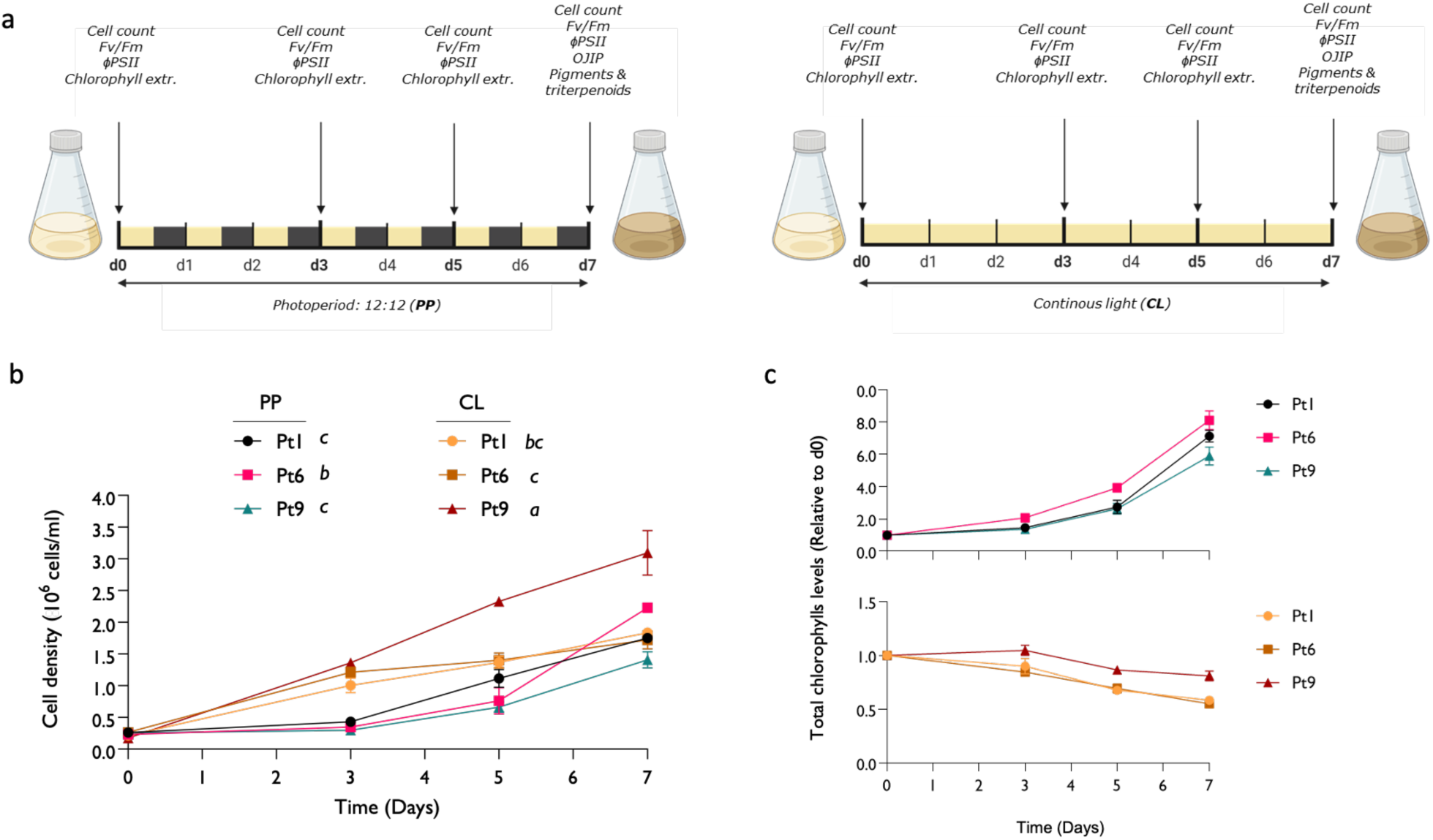
Effect of different light regimes on diatom cultures growth (a) Cartoon representing the experimental design of *P. tricornutum* cultures exposed to continuous illumination (CL) or 12 hrs:12 hrs light:dark photoperiod (PP). (b) Growth curve showing differences in cell density in cultures of Pt1, Pt6 and Pt9 grown in CL or PP. (c) Panels showing the change in total chlorophyll *a*/*c* content (compared to day 0 of culture) of cultures grown in either CL or PP. Plots show the mean and SD of 3 independent replicates. Different letters indicate statistically significant differences in the values reached at d7 (one-way ANOVA test, *P*<0.05).

### 2.3 Photosynthetic measurements

At specific time points during the experimental period (d0, d3, d5, and d7), 2 mL of each replicate culture was diluted with 2 mL of EASW to minimize signal saturation risk for bio-optical assessments. Chlorophyll-*a* pulse amplitude modulated (PAM) fluorometry (AquaPen-C, AP 110-C, Photon System Instruments, Brno, Czech Republic) was employed for bio-optical evaluation.

The effective quantum yield (ϕPSII) in light-adapted samples was calculated as ΔF/Fm’, where ΔF corresponds to Fm’-F (the maximum minus the minimum fluorescence of light-exposed algae). This measurement was conducted directly on samples obtained from the incubator, without any dark acclimation period, providing insight into the photosynthetic activity under growth light conditions. After this measurement, samples underwent a 20-minute acclimation in darkness. Chlorophyll transient light curves were then generated using the preprogrammed OJIP protocol, delineating fluorescence rise through four phases termed O, J, I, and P. [30]. Following the same protocol, the maximum quantum yield (Fv/Fm) was calculated as (Fm-Fo)/Fm, where Fm and Fo represent the maximum and minimum fluorescence of dark-adapted samples, respectively. Parameters derived from the curve and computed by the software from this analysis are detailed in Table 1.

**Table 1.** Summary of key parameters of the OJIP curve used to analyze photosynthetic efficiency and performance in photosystem II (PSII).

| Parameter | Description |
| --- | --- |
| Fv/Fm | Maximum quantum yield of PSII photochemistry ($Fv/Fm = (Fm - Fo) / Fm$) |
| ABS/RC | Apparent Light Energy Absorbed per Active PSII Reaction Center: Reflects the amount of light energy absorbed by each active PSII reaction center. Provides insights into the light-harvesting efficiency of PSII. A higher ABS/RC value suggests better light absorption capacity. |
| Plabs | Performance Index: Combines information from different phases of the OJIP curve. Quantifies the overall photosynthetic performance by considering both energy absorption and electron transport. Higher Plabs values indicate more efficient photosynthesis. |
| TRO/RC | Total trapped energy flux per active PSII reaction center. |
| Vi | Variable fluorescence intensity at the i-step (measured at 30 ms) in the OJIP curve. Provides information about the fluorescence yield during the transition from the dark-adapted state to the light-acclimated state. |
| Psi_o | Quantum yield of non-photochemical energy dissipation at the J-step. Calculated as $1 - Vj$, where $Vj$ is the fluorescence intensity at the J-step. Higher Psi_o values suggest increased energy dissipation pathways. |
| Phi_Eo | Quantum yield of electron transport beyond QA (from plastoquinone A to plastoquinone B). Calculated as $(1 - (Fo / Fm)) * Psi_o$. High Phi_Eo values indicate efficient electron flow beyond the initial QA reduction. |
| Vj | Variable fluorescence intensity at the J-step (measured at 2 ms) in the OJIP curve. Occurs shortly after the onset of illumination. $Vj$ reflects the reduction of plastoquinone A (QA), a key component in the photosystem II (PSII) electron transport chain. High $Vj$ values indicate efficient QA reduction and rapid electron flow through PSII. |
| ϕPSII | Effective quantum yield of PSII photochemistry in the light. Indicates the proportion of light absorbed by chlorophyll in PSII that is used in photochemistry. |

On the 7th day of growth under specific conditions, rapid light curves (RLCs) and NPQ kinetics were generated using preprogrammed LC1 and NPQ2 protocols, respectively. RLCs involved 6 incremental steps of actinic irradiance (E; 0, 10, 20, 50, 100, 300, 500 μmol photons m^-2^ s^-1^). At each step, ϕPSII was monitored every 60 seconds, and the relative electron transport rate (rETR) was calculated as E x ϕPSII.

The light response and associated parameters ETRm (maximum electron transport rate) and α (photosynthetic rate in the light-limited region of the light curve) were characterized by iteratively fitting the model of the rETR versus E curves using MS Excel Solver. [31]. The curve fitting was associated with r > 0.98 in all cases. Non-photochemical quenching (NPQ) was determined by tracking the progression of the Ft parameter (instantaneous chlorophyll fluorescence) while subjecting the samples to high light (HL, 500 μmol photons m-2 s-1) for 200 seconds, followed by a transition to very low light (LLRec, 10 μmol photons m-2 s-1) to facilitate NPQ relaxation. [11]. Fm (maximum fluorescence) was determined by administering a saturating pulse of light (SAT) every 20 seconds during HL and during LLRec. NPQ was calculated as (Fm-Fm’)/Fm’, where Fm denotes the maximum fluorescence in dark-adapted samples, and Fm’ represents the maximum fluorescence under subsequent conditions, computed via the SAT pulse.

### 2.4 Pigment extraction and analysis

Pigment extraction was performed as described by Mendes et al., 2007 [32]. Briefly, 3 ml of each diatom culture underwent centrifugation for 10 minutes at 4000 g and 21°C. The resulting pellet was rinsed with 1.5 ml of phosphate-buffered saline (PBS), followed by pellet collection through centrifugation under the same conditions. The collected pellet was flash-frozen in liquid nitrogen and stored at -80°C until further analysis. Samples were freeze-dried overnight, and pigments were extracted using 95% cold methanol buffered with 2% ammonium acetate. Extracts were filtered with 0.2 μm filters and 10 μl injected into an Agilent HPLC system through a YMC HPLC column 150 x 3.0 mm (S-3 µm) with a elution gradient of solvent A (40% acetone-60% methanol) and solvent B (40% water-60% acetone) as follows: 0-22 min from 0 to 30% B, 22-41 min from 30 to 10 % B and 45-55 min from 10 to 60% B. Column temperature was maintained at 40 °C while the flow rate was 0.5 mL·min^−1^. Pigments were identified via absorbance spectra and retention times by comparison with pure crystalline standards (DHI, Hørsholm, Denmark) To assess the activity of the xanthophyll cycle, samples were collected after each step of the NPQ protocol. The de-epoxidation state (DES) was calculated as Dtx/(Dtx+Ddx), where Dtx and Ddx represented diatoxanthin and diadinoxanthin, respectively.

Chlorophylls were quantified during the culturing period, through extraction and spectrophotometrical quantification as described in [33].

### 2.5 Triterpenoid analysis

Freeze-dried cells were lysed by incubation at 95°C for 10 minutes in equal volumes (250 μL each) of 40% KOH and 50% ethanol. Next, 900 μL of hexane was added, and the organic fraction was collected. This process was repeated twice, and the organic fractions were pooled and evaporated under nitrogen. The fractions were derivatized by adding 20 μL of pyridine and 100 μL of N-methyl-N-(trimethylsilyl)-trifluoroacetamide and by incubating the mixture for 1 hour at 70°C. GC-FID analysis was performed using an Agilent 7890B unit. 1 μL of sample was injected into a DB-5 column (30.0 m x 0.25 mm x 0.25 μm). The following settings were used: helium carrier gas was set at a constant flow of 1 mL/min; the oven temperature was programmed as follows: 80°C for 1-minute post-injection; ramped to 280°C at 20°C/min, held at 280°C for 45 minutes, ramped to 320°C at 20°C/min, held at 320°C for 1 minute, and cooled to 80°C at 50°C/min at the end of the run. Metabolite peaks were identified based on retention times of authentic standards (Sigma-Aldrich, USA). Peak area measurements were normalized by the weight of the dry matter.

### 2.6 Transcriptomic data analysis

Data used for this analysis came from [34] and were retrieved from the publicly available BioProjects PRJNA430316 and PRJNA971163. The quality of reads was assessed before and after trimming with Trim Galore [35] using FastQC [36] with default settings. Reads were mapped to the reference genome ASM15095v2 of *P. tricornutum* using HiSAT2 [37]. Quantitation was performed in SeqMonk [38]. Duplicates were removed, and default import settings for BAM files were used. Differential expression analysis was performed using DESeq2 after Benjamini and Hochberg correction. Due to substantial variation between replicates, the cumulative distribution of all samples was matched via “Matching Distributions”. Quantitation was conducted using the RNA-Seq pipeline quantitation, counting reads over CDS as raw counts, assuming a strand-specific library. Differential expression was considered significant with an adjusted p-value below 0.05. Differential expression in metabolic pathways, as defined by a previous study [39], were analyzed, and mosaic heatmaps were made using R Studio.

### 2.7 Cloning and diatom transformation by bacterial conjugation

#### 2.7.1 Transformation of *Phaeodactylum tricornutum* cells

The constructs *49202p_CrGES-mVenus_FcBPt*, cloned into the pPTPBR11 *E. coli* vector (Fabris et al., 2020), were used to transform Pt1, Pt6, and Pt9 via bacterial conjugation as described by Diner et al., (2016). Dry diatom discs were overlaid with 60 μl of bacterial culture for 1.5 hours in the dark, followed by 3 days under standard culture conditions. Transformed cells were then resuspended in 1 ml of ESAW, and 500 μl were plated in triplicates on ESAW 2% agar plates containing 100 μM Zeocin (Thermo Fisher Scientific, USA) for selection. The plates were incubated for 15 days at 21°C under continuous light (50 μmol photons m^−2^ s^−1^) until colonies were sufficiently large to be counted and picked.

#### 2.7.2 Flow cytometry analysis

To screen for the presence and expression of recombinant proteins, Pt1, Pt6, and Pt9 cells expressing *49202p_CrGES-mVenus_FcBPt*, were grown in 96-well round-bottom plates (Nunclon-treated, Thermo Fisher Scientific, USA) for four days. Screening was performed using a Guava easyCyte flow cytometer (Guava easyCyte HT, Cytek Biosciences, USA). mVenus fluorescence was measured using a 405-nm laser with the green spectral imaging band (525/40 nm). Cells were measured at a flow rate of 0.59 µl/s, with 5000 events per sample, in triplicates.

#### 2.7.3 Confocal microscopy

Cells expressing 49202p_CrGES-mVenus_FcBPt, at an OD750 of 0.2 were used for confocal microscopy analyses. Samples were imaged with an AX confocal/multiphoton system (Nikon, Japan) using a Plan Apo λ 100x oil-immersion objective with 1.49 NA (Nikon, Japan) at a resolution of 1024 × 512 pixels. mVenus was excited with a 488 nm argon laser and chlorophyll autofluorescence with a 561 nm laser. The fluorescence emission of mVenus was detected at 500-520 nm, and chlorophyll at 625-720 nm. Light was passed through the transmitted light channel. Images were processed using ImageJ software.

### 2.8 Geraniol capture and quantification

Geraniol was captured by adding 2 mL of isopropyl myristate (IM; Sigma-Aldrich, USA) to 50 mL of diatom culture at the time of inoculation. In all experiments, the IM layer was harvested by centrifugation for 10 min at 4500g at the end of the cultivation (13 days) and directly analyzed by GC-MS. Samples were manually injected and run on an Agilent 5977C GC/MSD system. The column used was an HP-5 (5%-phenyl)-methylpolysiloxane nonpolar column (30.0 m × 0.32 mm × 0.25 μm). The carrier gas was helium, used at a constant flow of 1.0 mL/min and an injection volume of 1 μL. The temperature gradient of the oven was 35 °C for 3 min, then sequentially increased at the rate of 20 °C per minute to 320 °C and then maintained for 5 minutes. The mass spectrometer operated in electron impact mode at 70 eV. The injector temperature was maintained at 280 °C and ion source temperature at 230 °C. Quantitative analysis of geraniol was run in Selective Ion Monitoring (SIM) mode, where selected ions were monitored for 20 ms each. The ion monitored was m/z 69.1. The peak areas were converted into metabolite concentrations with calibration curves plotted against a range of known concentrations of geraniol standard (Sigma-Aldrich, USA).

## 3 Results and discussion

### 3.1 Influence of light regime on the growth of three different *P. tricornutum* strains

*P. tricornutum*, though not highly abundant, is cosmopolitan and thrives in diverse locations globally. Its resilience is evident in its adaptation to fluctuating environments like estuaries and rock pools (De Martino et al., 2007). Variability in water depths, affecting light availability, is crucial for diatom growth. We evaluated the influence of light regimes comparing photoperiodic (PP, 12 hours light/12 hours dark) versus continuous illumination (CL) (Fig.1a), on the growth of three *P. tricornutum* strains (Pt1, Pt6, and Pt9) in a defined medium under atmospheric CO2. During PP growth, each of the three strains demonstrated a distinct initial lag phase lasting approximately three days, a period during which no significant changes in cell density were observed across the strains. Following this phase, the exponential growth commenced on the fourth-day post-inoculation and was sustained throughout the seven-day duration of the experiment. Strain Pt6 showed substantially higher growth, with its cell density increasing eight-fold from the initial value by day seven. The growth of this strain was more pronounced than that of strains Pt1 and Pt9, which both exhibited a six-fold increase in OD. Notably, the faster growth of Pt6 was particularly evident during the latter three days (days 5 to 7) of the study, as evidenced by the steeper slope when compared to Pt1 and Pt9 (Fig. 1b).

Whereas strains Pt6 and Pt1 showed similar growth under CL, each increasing their cell density seven-fold, strain Pt9 had the most robust growth with a ten-fold cell density increase. Overall, all three strains exhibited faster growth when subjected to continuous light, although their growth responses varied between the two lighting conditions (Fig. 1b). This variation may be partially attributable to the natural habitats of the different strains, which could influence their adaptive responses to different light regimes. Certain microalgae species are particularly well-adapted to fluctuating light and dark cycles, as the dark periods allow for the repair of their photosynthetic apparatus from prolonged and excessive light exposure, thereby supporting sustained growth [41]. In contrast, other species thrive under continuous light conditions by maintaining constant photosynthetic activity, which can lead to faster growth rates, provided they can avoid photoinhibition [42].

Pt6 was isolated from intertidal rock pools. Here, the environmental conditions including temperature, salinity, wave action, and rapid changes in light irradiance, would favor light-dark alternation traits, as many intertidal microalgae have developed circadian rhythms that synchronize their metabolic activities with tidal cycles (Lewin et al., 1958; Dolganyuk et al., 2020), In contrast, the tropical Pt9 strain thrives under CL, maximizing photosynthesis and carbon fixation without photoinhibition, resulting in an improved reproduction rate as observed in various tropical marine phytoplankton exposed to prolonged illumination (Brand and Guillard, 1981).

The growth pattern of different diatom strains correlates with their chlorophyll *a* content. Under alternating light and dark conditions, this was similar among the three strains, increasing with culture density. However, CL caused variations in pigment accumulation as chlorophyll levels decreased in all strains but more slowly in Pt9 compared to Pt1 and Pt6 (Fig. 1c). CL leads to excess photons similarly to what happens during exposure to high irradiance, causing a decrease of the chlorophyll molecules composing the light harvesting complexes to prevent oxidative damage to photosystems, a response observed in *P. tricornutum* and the green alga *Chlamydomonas reinhardtii* [39,40,41,42]. The tropical origin of P9 likely makes this strain more resilient in CL conditions, maintaining higher chlorophyll levels and better tolerating oxidative stress without significant chlorophyll reduction, as tropical diatoms typically withstand higher irradiance and temperature [48].

### 3.2 Effect of light regimes on the photo-physiology of the three *P. tricornutum* strains

Considering the effect of different light regimes on the growth and chlorophyll content of Pt1, Pt6, and Pt9 strains, we hypothesized that varying light exposure would also influence their photosynthetic efficiency. Variations in the photosynthesis of *P. tricornutum* strains have been previously documented [34], however, no studies have yet investigated the impact of varying light conditions on the cultivation of different strains. Thus, we measured the maximum quantum yield (Fv/Fm) and effective quantum yield (ΦPSII) under different light conditions to assess the potential maximal and actual photosynthetic activity of diatoms during growth. Fv/Fm indicates the maximum efficiency of photosystem II under dark-adapted conditions and is used to detect stress before visible symptoms such as culture bleaching and delayed growth (Kramer et al., 2004). Optimal values for C3 plants are around 0.83–0.84, but are generally lower in algae and lichens, and decrease with photoinhibition [49]. ΦPSII, typically lower than Fv/Fm, measures actual photosynthetic efficiency during active growth and is highly sensitive to environmental changes, offering real-time insights into photosynthetic behavior under stress [49]. Similarly to growth data, both the Fv/Fm and ΦPSII remained stable throughout the PP culture period **(Figure 2a, b).**

**Figure 2:**
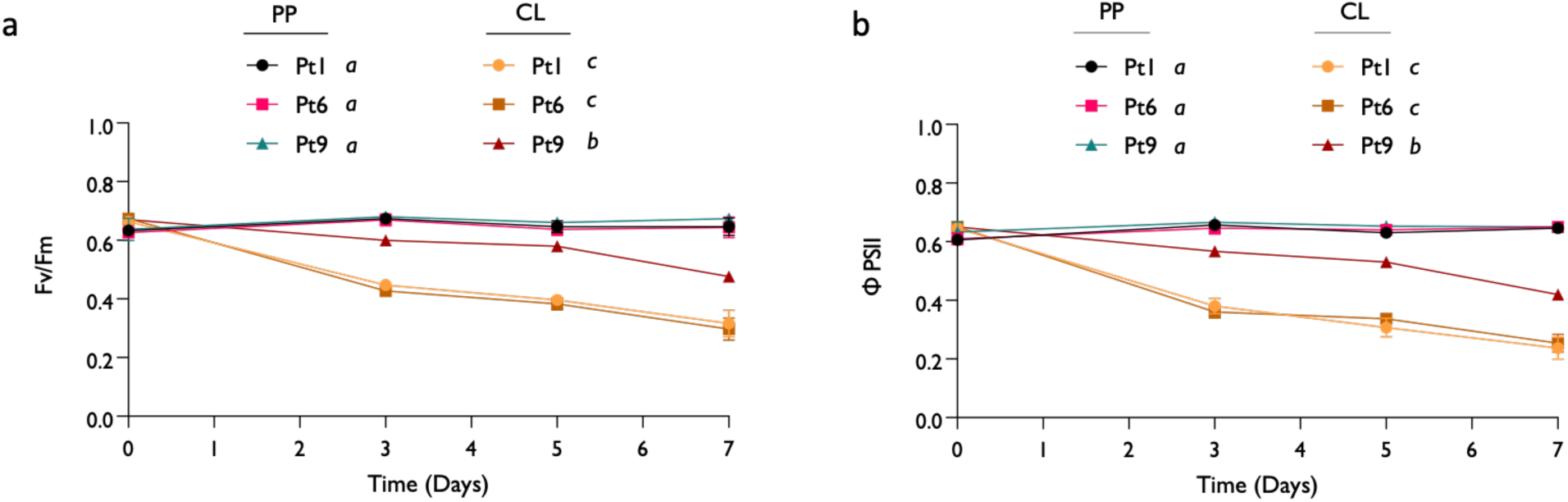
Impact of different light regimes on the photosynthetic activity of different *P. tricornutum* strains. The panels show (a) the changes in maximum quantum yield (Fv/Fm) and (b) effective quantum yield (ϕPSII) in the three strains grown under different light regimes. Plots show the mean and SD of 3 independent replicates. Different letters indicate statistically significant differences in the values reached at day 7 (one-way ANOVA test, *P*<0.05).

These findings indicate that this light regimen did not exert significant stress on the photosynthetic machinery, thereby not impacting photosynthetic performance adversely. Differently, when cultivated under continuous light, the photosynthetic efficiency decreases after 3 days of culture. Although both regimes deliver the same light intensity, CL has been demonstrated to induce a photo-inhibitory effect akin to that caused by high light in plant models [50]. In high light-tolerant algae like *Nannochloropsis*, optimal growth involves alternating light and dark cycles. In fact, dark periods are crucial for re-oxidizing electron transporters [51] in the photosynthetic apparatus and without photoperiods, algae can suffer radiation damage, significantly reducing photosynthetic productivity [52].

In our experimental set-up, the continuous illumination could induce excessive pressure on the photosynthetic apparatus, associated with oxidative stress that would progressively reduce the overall health of the culture. Among the three strains, Pt9 demonstrated greater tolerance to the light treatment, exhibiting less decline in photosynthetic performance (Fig. 2). This aligns with observations on the growth performance, suggesting that this strain may possess innate mechanisms for protection against light stress.

To characterize specific aspects of the photosynthetic performance of the three strains, we performed several chlorophyll *a* variable fluorescence measurements on the 7^th^ day of growth in the respective light conditions. We started from a OJIP test as an analysis of the fate of photons absorbed by the PSII antennae (trapping, forward electron transport beyond QA and dissipation as heat) [53]. In algae and higher plants, under optimal conditions and short dark adaptation, PSII centers are open with a fully oxidized primary electron acceptor QA, resulting in minimal chlorophyll fluorescence yield (Fo). Under strong illumination all PSII centers are closed, and maximum fluorescence yield (Fm) is achieved. The OJIP kinetics of light-induced chlorophyll fluorescence in dark-adapted green algae and plants show several inflection points, with the minimum O (F50μs) and maximum P (Fp) fluorescence signals corresponding to Fo and Fm, respectively [54]. The rapid OJ rise is associated with the reduction of QA, whereas the following JIP phase reflects the further reduction of QA due to the decrease in the rate of its re-oxidation, which is modulated mainly by the redox state of plastoquinone (PQ) pool [55]. Typically, the shape of the initial fluorescence rise depends on the energetic connectivity between PS II units and in organisms cultivated under optimal conditions the shape of the initial fluorescence rise shows sigmoidicity, associated with the high probability of exciton exchange between PSII units in a supercomplex [56].

Under PP, the three strains displayed the characteristic shape of the OJIP curve (Fig. 3a). However, curves from cultures grown in CL were typically flatter, indicating that this condition may not be optimal. Pt9 uniquely showed a tendency towards a sinusoidal shape in the OJIP kinetics, reinforcing our findings that this strain is more adapted to continuous light conditions (Figs. 1-2). Consistently, Pt9 also exhibits the lowest value of ABS/RC, a parameter indicative of changes in PSII antenna size. This could result from alterations in the number of LHC complexes per PSII reaction center or the inactivation of reaction centers [57]. Furthermore, a smaller antenna size correlates with fewer inactivated reaction centers and a higher performance index (PIabs), as observed in Pt9.

**Figure 3:**
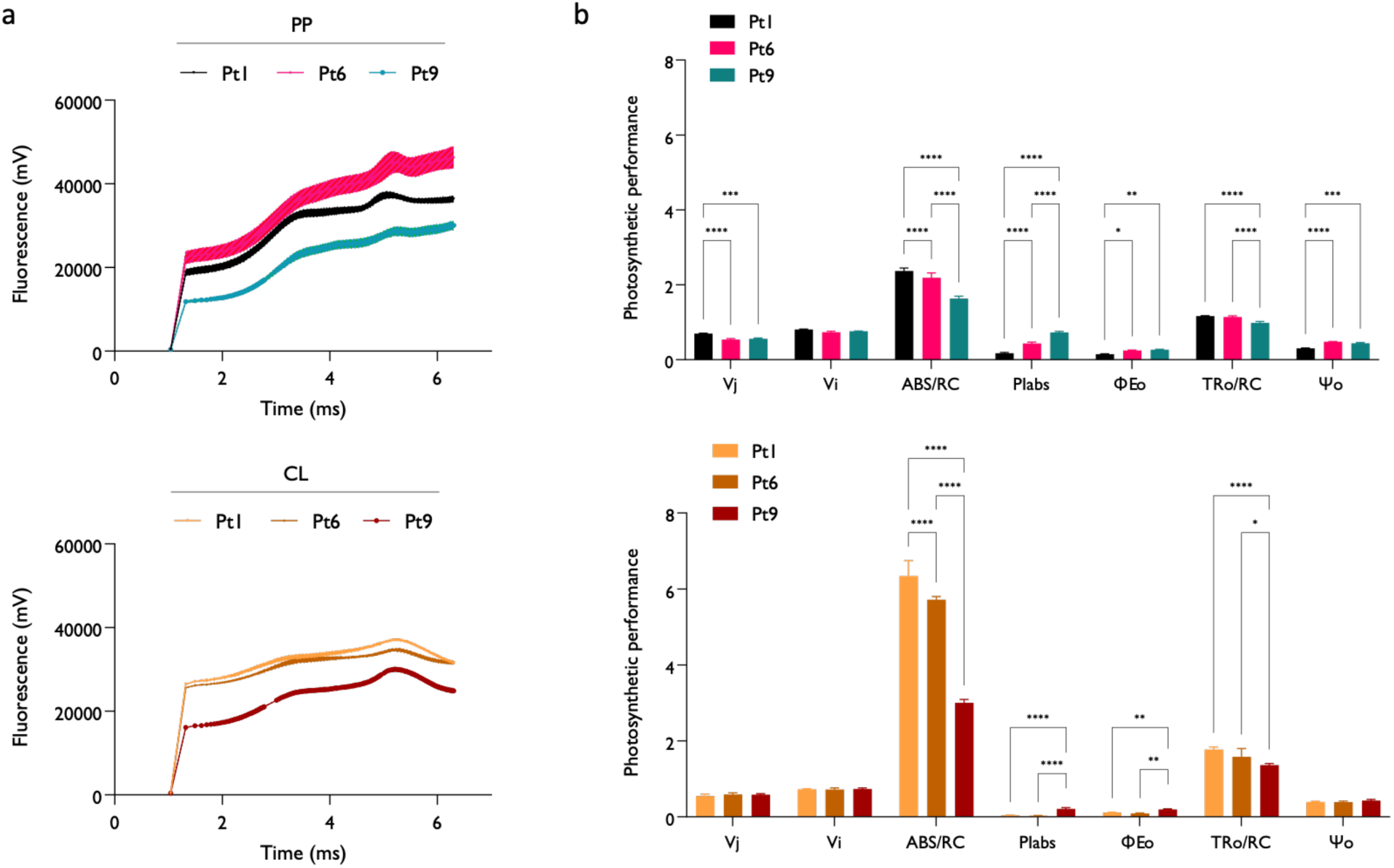
Effect of growth light regime on the dynamics of electron circulation in PSII. (a) OJIP curves for Pt1, Pt6, Pt9 grown under PP or CL for 7 days; (b) OJIP curve-derived parameters (abbreviations explained in Table 1). Plots show the mean and SD of 3 independent replicates. Asterisks indicate statistically significant differences (Two-way ANOVA, *: *P*<0.01, **: *P*<0.05, ***: *P*<0.001, ****: *P*<0.005).

At the end of 7 days of cultivation in different light regimes we further investigated the photosynthetic behavior of Pt1, Pt6 and Pt9, by calculating rapid light curves (RLCs) to assess the PSII functionality during the exposure to different incremental light intensities [22].

RLC are generally higher in samples grown under a 12hrs:12hrs photoperiod, indicating this might be the optimal condition for all three strains of *P. tricornutum*. In all samples, the maximum relative electron transport rate (rETRm) was higher when cultivated under photoperiod and lower in continuous light, likely due to excessive stress on the photosynthetic apparatus (Fig. 4b).

**Figure 4:**
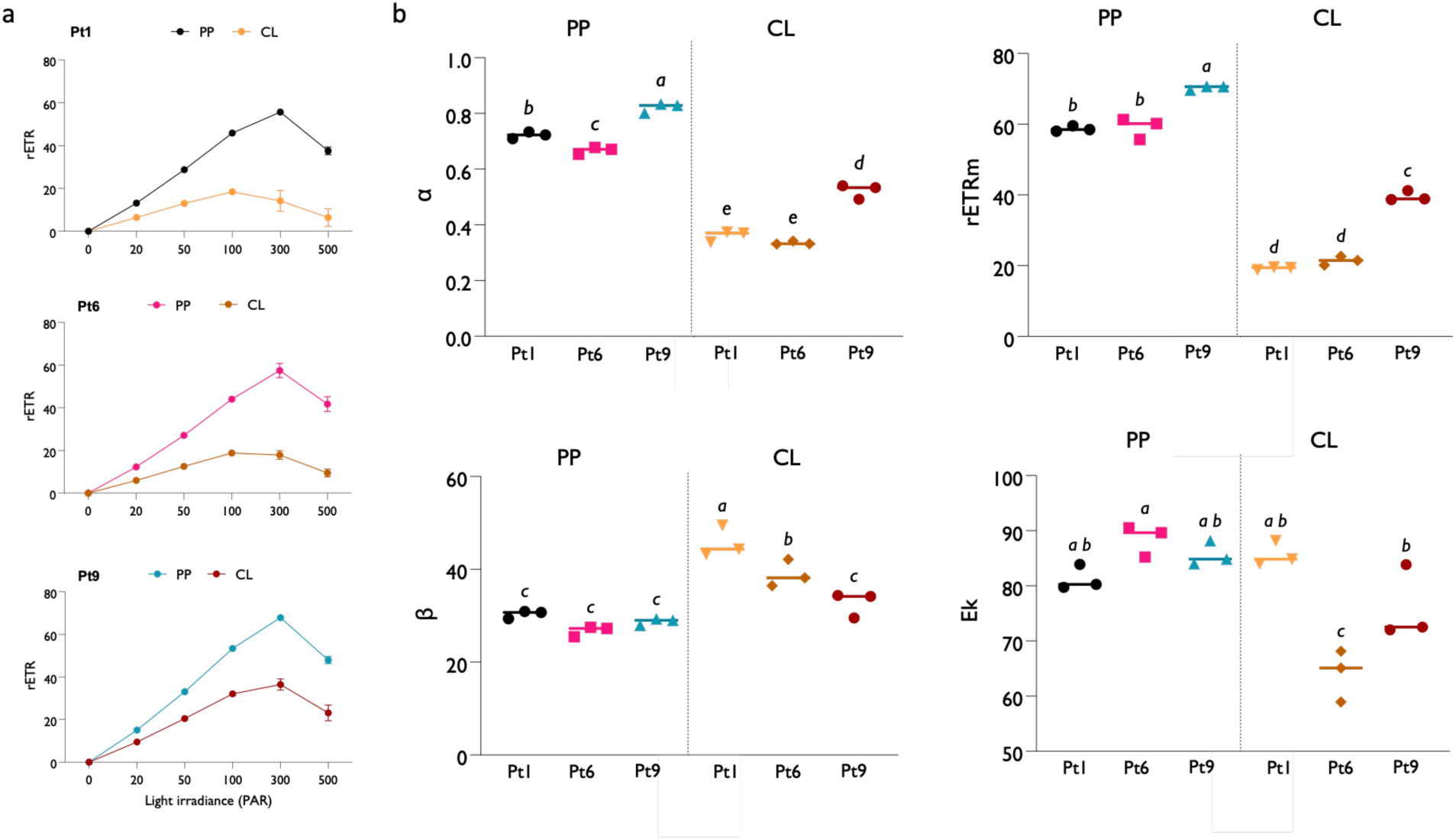
Photosynthetic response of the three strains exposed to increasing light intensities. (a) RLCs calculated for Pt1, Pt6 and Pt9 grown under different light regimes for 7 days. The right panels (b) show the curve kinetic parameters calculated from the RLC using the Platt model. Plots show the mean and SD of 3 independent replicates. Different letters indicate statistically significant differences in the values (one-way ANOVA, *P*<0.05).

Notably, among the three strains, Pt9 consistently outperformed the others. This, not only achieved a considerably higher ETR but also exhibited a higher photosynthetic rate in the non-light-limiting part of the curve (α) and a lower coefficient of photoinhibition (β), alongside a higher light saturation coefficient (Ek) [58]. Thus, although CL is not optimal, Pt9 diatoms cultivated under these conditions have superior photosynthetic performance across a broader range of light intensities, thus displaying significant adaptability and suitability to environments with abundant light, such as photobioreactors or photosynthetic incubators. As a further confirmation, we profiled the photoprotective capacity of the three strains by calculating their level of non-photochemical quenching (NPQ) by exposing dark-acclimated samples to short-term high light stress and subsequently returning them to dim light to assess the recovery of photosynthetic functions.

Non-photochemical quenching (NPQ) allows diatoms to prevent photodamage to the photosynthetic apparatus from excessive light energy. This process involves the dissipation of excessively absorbed light energy as heat through various xanthophyll pigments. In the case of *P. tricornutum*, this involves the dynamic conversion of diadinoxanthin (Ddx) to diatoxanthin (Dtx) upon exposure to high irradiance [41].

NPQ in diatoms is dynamic, adjusting quickly to changes in light intensity, which is essential for their survival in fluctuating light environments typical of aquatic habitats where they often reside [21,59]. PP-grown Pt6 and Pt1 both exhibited a high level of non-photochemical quenching (NPQ) build-up during exposure to high light, which was almost completely dissipated when the samples were returned to low irradiance. Conversely, Pt9 developed a significantly lower NPQ level under high light, which similarly dissipated in low light (Fig. 5a). Thus, Pt1 and Pt6 are more reactive to high light stress, whereas Pt9 does not exhibit the same level of stress response.

**Figure 5:**
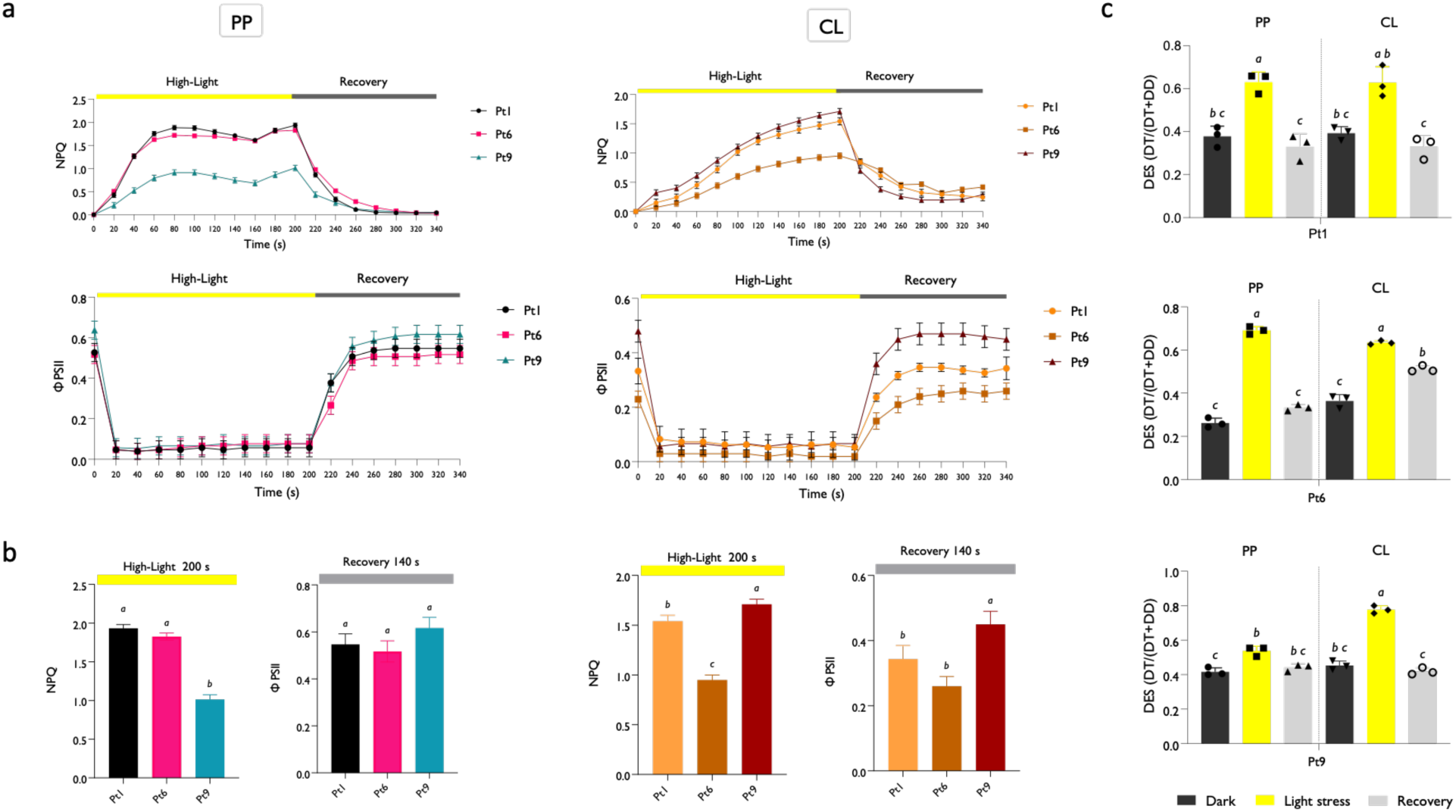
Variable response to short-term high light stress in the three strains of *P. tricornutum*. (a) The upper panels show the NPQ value in PP-grown samples exposed to high light (yellow box, 500 PAR) for 200 s and subsequently exposed to a recovery period under dim light (20 PAR) for 240 s. The lower panels show the value of effective quantum yield in the same conditions. (b) The lowest bar plots correspond to the NPQ and ϕPSII values observed after 200 s in high light and after 240 s in recovery, respectively. The two sets of graphs show the data for samples exposed to PP and CL; (c) De-epoxidation state calculated in the same samples after dak acclimation (dark grey bar), high light stress (yellow bar) and recovery (light grey bar). Plots show the mean and SD of 3 independent replicas. Different letters indicate statistically significant differences (one-way ANOVA, *P*<0.05).

The conversion of Ddx to Dtx defined by the De-epoxidation State (DES) value (Fig. 5c), reflects this kinetic response. Under both photoperiod and continuous irradiance conditions, Pt1 and Pt6 accumulate high levels of diatoxanthin. Notably, Pt6 builds up the highest DES value among the three strains, while the levels remain nearly unchanged in PP-cultivated Pt9. This suggests a differential endogenous carotenoid level, likely supported by a strain-specific gene expression variability, impacting the diadinoxanthin and diatoxanthin ratios, even during short-term light stress exposure [60]. In continuous light conditions (Fig. 5b), the dynamics shift, with Pt9 exhibiting the highest NPQ levels among the strains and the greatest capacity to recover photosynthetic activity at lower irradiance. This aligns with our previous observations on the capacity of Pt9 to maintain high NPQ levels, which are consistent with its geographic distribution in latitudes that receive intense solar radiation [15,56]. Conversely, Pt6 shows a reduced ability to lower the DES value after exposure to reduced irradiance, suggesting a potentially lower photoprotective capacity, which relies on the thylakoidal proton gradient [61].

### 3.3 Effect of light regimes on pigment composition in *P. tricornutum* strains

Based on our observations of the photoprotective abilities of the three strains, we hypothesized that the different strains might respond to light regimes with varying pigment compositions and that this variation could result from differences in the levels of terpenoid precursors. Moreover, a differential pigment composition could have potential biotechnological applications due to the industrial relevance of carotenoids. To verify this, we analyzed the content of pigments, distinguishing between photoprotective (diadinoxanthin, diatoxanthin, and β-carotene) and light-harvesting ones (chlorophyll a, c, and fucoxanthin) [62]. In accordance with previous observations (Figs. 1, 5), the total pigment levels were higher in PP-cultivated strains (Fig. 6, Fig. S1).

**Figure 6:**
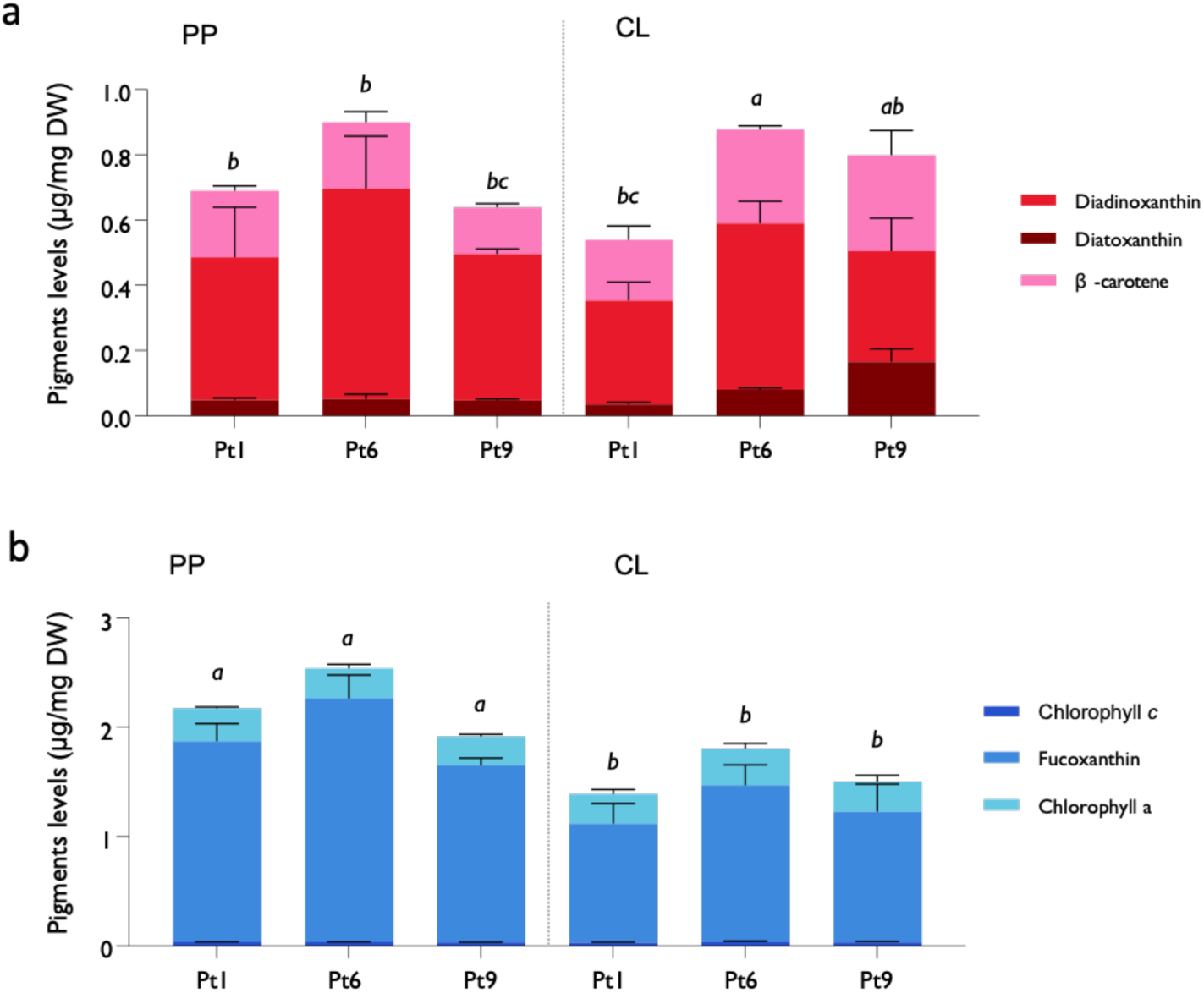
Pigments content of Pt1, Pt6 and Pt9 exposed to different light regimes. The upper graph (a) shows the accumulation of carotenoids with a photoprotective role, while the lower panel (b) shows that of pigments with photon capture function. Pigments levels are expressed as μg/mg of dry weight and calculated with a calibration curve of the authentic standards. Plots show the mean and SD of 3 independent replicates. Different letters indicate statistically significant differences in the levels of total pigments between samples (one-way ANOVA, *P*<0.05).

The observed differences mainly regarded the increased levels of light-harvesting pigments (e.g., fucoxanthin, chlorophyll a, and chlorophyll c), which accumulate during the light-dark cycle and are likely necessary to maximize photon capture—an essential process not as critical under continuous light conditions [63]. Pt6 exhibited higher levels of fucoxanthin compared to Pt1 and Pt9, and correlates with the photosynthetic data which indicated that Pt6 was the best performer under variable light intensity **(Fig. S1).** In this scenario, it is plausible to assume that its photon-capturing ability, enhanced by fucoxanthin, is superior. Moreover, this strain was already identified as optimal for fucoxanthin enrichment due to its higher expression of the violaxanthin de-epoxidase-like protein 1 gene (VDL1), which catalyzes the conversion of violaxanthin to neoxanthin in the fucoxanthin biosynthesis pathway, compared to other strains when cultivated under continuous low light (60 μmol photons/(m²·s) [64]

Under CL, on the other hand, the content of light-harvesting pigments decreased because, in conditions of light oversaturation, diatoms reduce the synthesis and accumulation of these pigments to mitigate excessive energy harvesting and prevent photodamage or photoinhibition [65] as, even at moderate intensities, prolonged light exposure can simulate the effects typically seen under high light conditions [66]. Notably, the Pt9 strain, which demonstrated increased tolerance to continuous light stress in this study, likely owes this to the highest basal diatoxanthin content among the strains. This could be attributed to a correlation between violaxanthin epoxidase gene activity, diatoxanthin production, and non-photochemical quenching (NPQ) response to high light which have been well-documented [67]. As with Pt6, the naturally high level of diatoxanthin in Pt9 would make it an optimal candidate for producing high yields of this compound, a feat previously achieved only through genetic engineering [68].

### 3.4 Effect of light regimes on triterpenoid composition in different strains

The accumulation capacity and composition of endogenous isoprenoid-derived molecules such as triterpenoids, are primary parameters to consider when designing metabolic engineering strategies aimed the production of endogenous or heterologous terpenoid-based high-value products. Hence, in addition to carotenoids, we set out to investigate whether different strains exhibit different triterpenoid compositions under various light regimes. Carotenoids and sterols are the main terpenoid sinks in *P. tricornutum* [26] and they are synthesized mainly from precursors produced by the methyl-erythritol phosphate (MEP) and mevalonate (MVA) pathways, respectively. Differences in the accumulation of these end-products are informative to understand the capacity and dynamics of the metabolic fluxes through these pathways in different strains and conditions. In particular, we focused on the main sterol compounds synthesized and accumulated by *P. tricornutum*: brassicasterol, campesterol and cholesterol, and their precursor squalene [69,70]. Under continuous light cultivation, the total sterol content was significantly higher in all three strains compared to growth with an alternation of light and darkness (Figure 7a). As previously observed, longer photoperiods generally accelerate growth, thereby promoting the accumulation of compounds such as sterols as the diatoms progress more rapidly through their growth phases, each associated with a specific level of sterol accumulation [71]. The variation in available light predominantly affects minor sterols such as campesterol, cholesterol, and their precursor, squalene **(Fig. 7, Fig. S2).**

**Figure 7:**
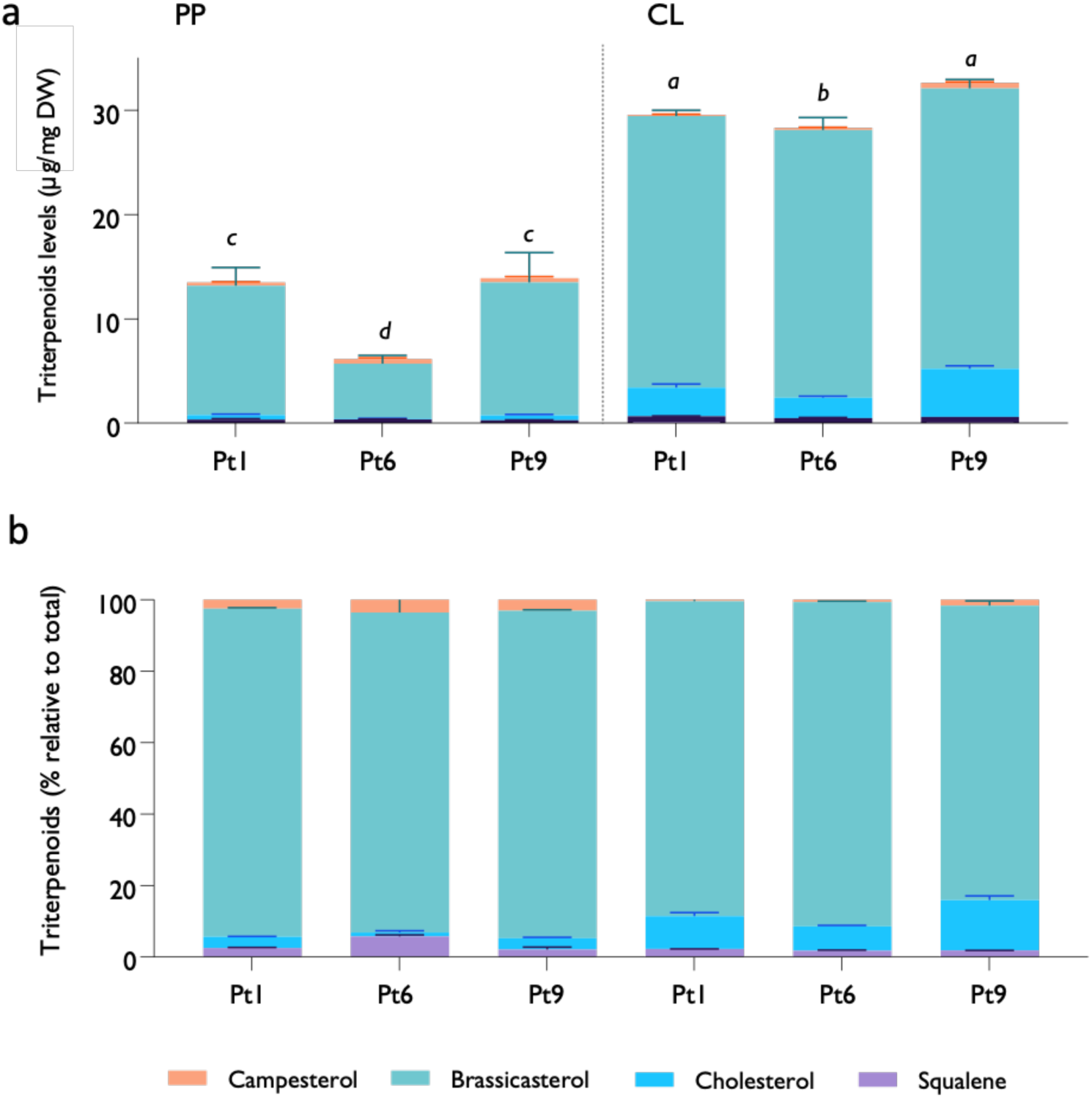
Triterpenoids composition of Pt1, Pt6 and Pt9 exposed to different light regimes. The upper graph (a) shows the absolute levels of triterpenoids as μg/mg of dry weight and calculated with a calibration curve of the authentic standards. The lower plot represents the relative percentage of the single triterpenoids over the total amount. Plots show the mean and SD of 3 independent replicates. Different letters indicate statistically significant differences in the levels of total pigments between samples (one-way ANOVA, *P*<0.05).

The metabolic shift in response to continuous light [72] correlates with reduced squalene accumulation in CL-grown samples, suggesting a preferential conversion of this into downstream products. Indeed, changes in culture conditions such as temperature and salinity have been shown to affect sterol composition, and light exposure likely exerts a similar impact [73]. Under CL exposure, the increased levels of cholesterol may aid in stabilizing internal membranes, which also relates to the variations in photosynthetic efficiency previously mentioned and the high-light stress response observed under CL conditions. Furthermore, in other algae such as *Haematococcus pluvialis*, high light stress has been associated with the simultaneous synthesis of sterols and triacylglycerides (TAG), leading to increased cholesterol production [74].

Therefore, it is possible to delineate a link between growth and photosynthetic performance and the inherent ability of different *P. tricornutum* strains to accumulate compounds of interest, revealing considerable genetic diversity. This diversity results in a robust metabolism linked to distinct classes of compounds, and understanding this genetic variation is key to selecting the best strain for biotechnological applications.

### 3.5 Transcriptional basis of the physiological and metaboilic differences among strains

Our previous results indicated that Pt1, Pt6, and Pt9 present differences at the physiological and metabolic levels, suggesting potential basal variations at the transcriptomic level supporting their metabolic plasticity. To provide a transcriptional context to the observed phenotypes, we mined the transcriptomics dataset generated by Chaumier et al. [34] obtained from samples cultivated at 19°C under a 12:12 light-dark cycle with a light intensity of 70 μmol/m²s. Principal component analysis (PCA) revealed significant variance, especially among Pt1 duplicates with R-squared values between duplicates of Pt1, Pt6, and Pt9 being 0.76, 0.8, and 0.99, respectively (Fig. S3). This indicated a substantial variation, particularly in Pt1 which limited the detection of differentially expressed genes (DEGs) in Pt1 compared to Pt6 and Pt9, with 124 and 742 DEGs, respectively (p-value < 0.05). DEG analysis between Pt9 and Pt6 revealed instead a higher number of DEGs (2925, p < 0.05), leading to a focus on this comparison.

Gene Ontology (GO) analysis of DEGs between Pt9 and Pt6 showed that downregulated genes in Pt9 were linked to chlorophyll binding, photosynthesis, nitrogen regulation, pigment biosynthesis, and thylakoid-associated processes [75]. Analysis of selected metabolic pathways suggested lower enrichment of genes involved in the Calvin-Benson cycle, PSI components, light-harvesting proteins, and chlorophyll biosynthesis in Pt9 compared to Pt6. Terpenoid metabolism, including the MVA pathway, sterol, and pigment biosynthesis, also showed downregulation (Fig. 8).

**Figure 8:**
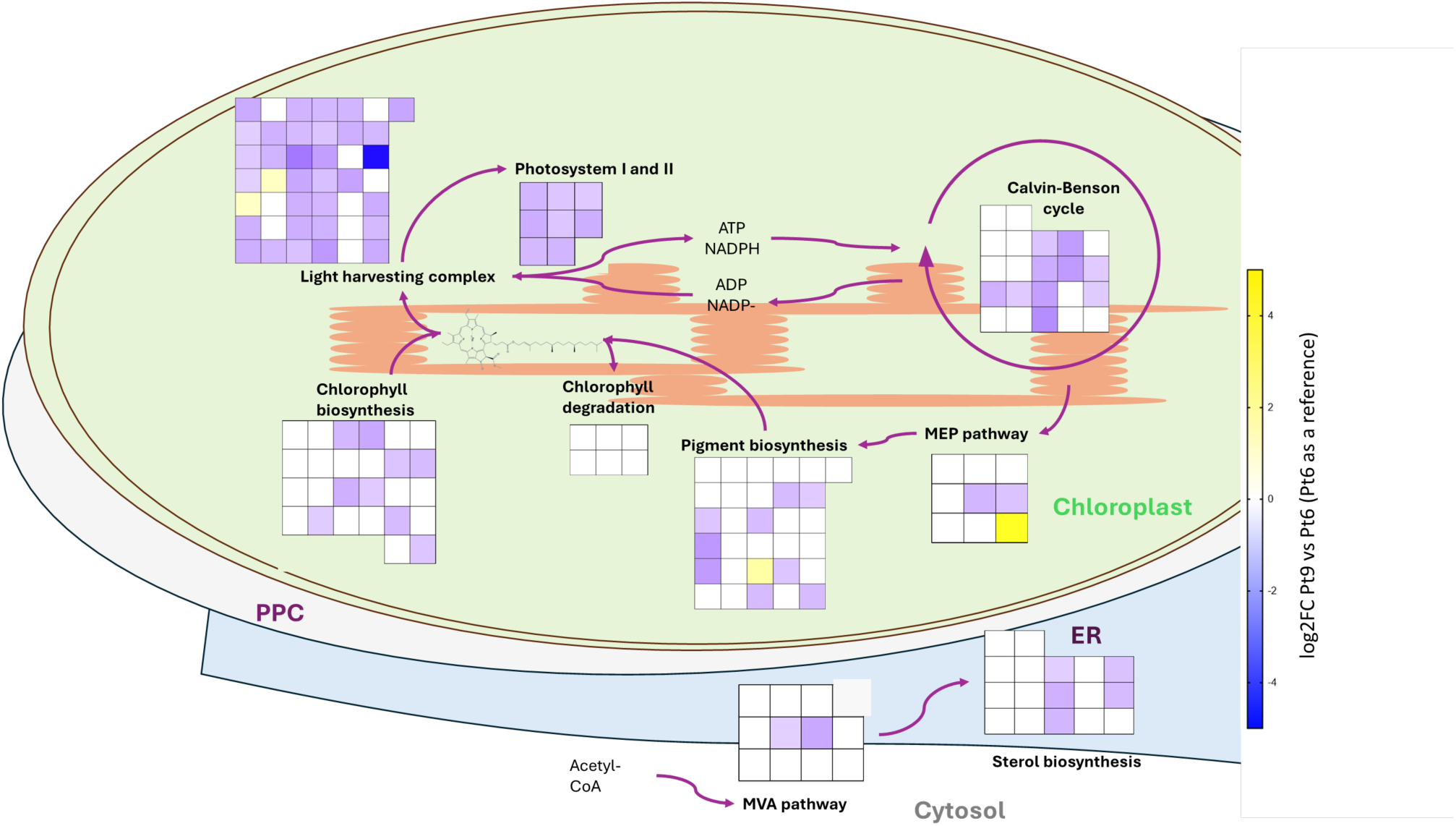
Schematic overview of metabolic pathways involved in photosynthesis and isoprenoid metabolism in *P. tricornutum*. Heatmaps representing GO term enrichment values (Z-scores) calculated from log2FC values from [34] comparing Pt9 and Pt6 (using Pt6 as a reference) genotypes. Only GO terms which were determined to be relevant for the physiologic and phenotypic variations determined in this work are shown. Detailed gene-annotation lists, organized by metabolic pathway are available in Supplementary Data File 1.

The higher growth rate of Pt6 in the exponential phase correlates with relatively higher expression of genes involved in photosynthesis, carbon fixation, and pigment synthesis. Conversely, the downregulation of these genes in Pt9 compared to Pt6 correlates with our observations of reduced photosynthetic activity, slower growth rate, and lower pigment content in Pt9 under a 12:12 light regime (Fig. 1). This is likely due to Pt9 being isolated from a tropical climate, where it prefers long-day cultivation conditions, unlike the 12:12 light: dark regime used in [34]. The downregulation of sterol biosynthesis genes in Pt9 compared to Pt6 contrasts with the higher sterol accumulation observed in Pt9 in this study (Fig. 7). This discrepancy may be due to post-translation regulation or negative feedback mechanisms, which, although not demonstrated in *P. tricornutum*, are common in animals, plants, and fungi [72,73,74,75].

Overall, the differences in pathway regulation between Pt6 and Pt9 are noteworthy, especially considering the strains were isolated from distinct environments but grown under identical conditions for transcriptomic comparison. These variations might favor the growth of one strain over the other in varying conditions, explaining the observed changes in photosynthesis-related gene expression and growth rates.

### 3.6 Conjugation efficiency and potential for monoterpenoid engineering

Considering the observed metabolic differences, we hypothesized that the three strains may have distinctive potential for metabolic engineering, for the heterologous production of high-value compounds, such as monoterpenoids and derivatives [26]. We previously demonstrated that diatoms like *P. tricornutum* might hold the potential to be engineered to synthesize these high-value compounds and possibly their derivatives, since they naturally accumulate free pools of the precursor geranyl diphosphate (GPP), differently from most yeast and bacterial production hosts [24,76]. A considerable degree of diversity – yet uncharted - is expected to exist in diatoms concerning their natural and engineered capacity to produce secondary metabolites. For example, while Pt1 has been shown not to produce any monoterpenoids naturally [26], it has been reported that a *P. tricornutum* strain isolated from Western Norwegian Fjord Water was found to produce volatile organic compounds (VOCs) with bioactive action, including monoterpenoids [81]. In this perspective, a strain that is naturally predisposed to synthesize this class of compounds might be highly attractive for designing and engineering more efficient strains.

For this reason, we investigated the natural production of monoterpenoids in Pt6, and Pt9 strains grown in continuous light (CL) or photoperiod (PP) with gas chromatography-mass spectrometry (GC-MS). However, none of the strains showed any monoterpenoid emission, indicating they do not naturally produce these compounds, at least in the cultivation settings used (Fig. S4)

Next, we investigated whether the three *P. tricornutum* strains would respond differently to genetic transformation via episomal DNA introduced by bacterial conjugation and to produce heterologous monoterpenoids, using geraniol as a model product. Episomes and bacterial conjugation are emerging new ways to generate transgenic diatom cell lines [40,82], they offer a reliable, consistent, and predictable platform for protein expression, with features that enable advanced synthetic biology approaches [80,80,83]. This approach avoids complications associated with random chromosomal integration, such as multiple insertions, position-specific effects on expression, and potential knockout of non-targeted genes [7]. Significant differences were observed among the three strains upon transformation with the *pPTBR11_Phatr3_J49202p_CrGES-mVenus* episome, carrying a gene encoding the geraniol synthase of *Catharanthus roseus* (CrGES) fused at its carboxy-terminus to an mVenus Yellow Fluorescent Protein (YFP), and controlled by the constitutive promoter region of the gene *Phatr3_J49202* [26]. In two independent conjugation events, Pt6 produced a significantly higher number of colonies compared to Pt1 and Pt9. However, none of these colonies expressed the fluorescent fusion protein, as determined by flow cytometry screening (Fig. 9a). In fact, Pt1 and Pt9 exhibited high mVenus fluorescence intensity, while Pt6 showed negligible fluorescence, attributable only to the autofluorescence of chlorophyll, which fell below the background level (Fig.S5). Additionally, confocal microscopy confirmed the presence of the green fluorescence signal only in the same strains (Fig. 9b), indicating that only Pt9 and Pt1 - as previously reported by [26] - are capable of expressing and correctly folding the *CrGES-mVenus* construct.

**Figure 9:**
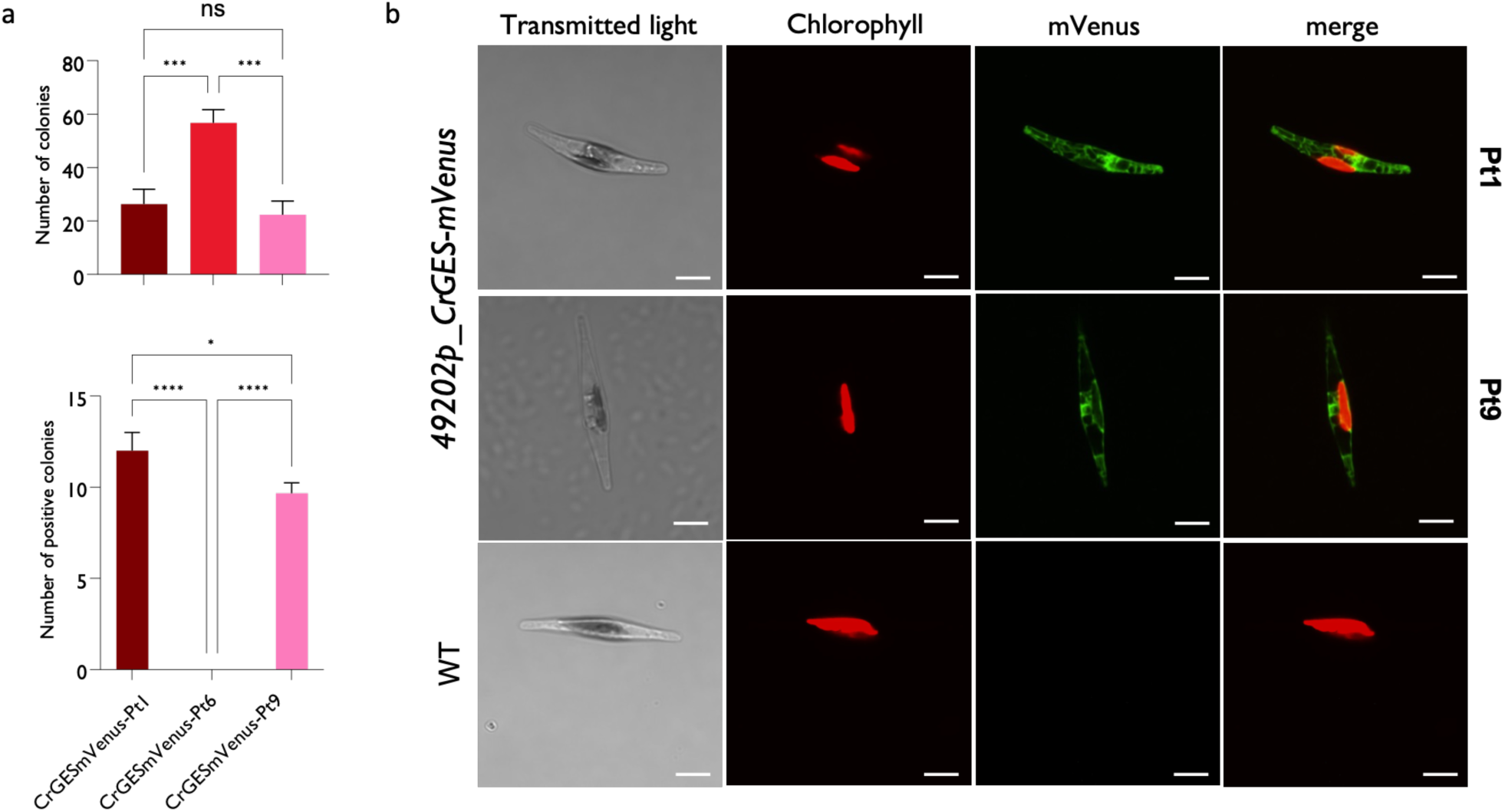
(a) Conjugation efficiencies in different *P. tricornutum* strains, determined based on the number of colonies obtained after two independent transformation events in the same conditions. The lower panels represent the number of colonies exhibiting mVenus fluorescence upon expression of the CrGES-mVenus fusion protein, quantified by flow cytometry. Asterisks represent significant differences (Two-way ANOVA, *: *P*<0.01, ***: *P*<0.05, ****: *P*<0.005); (b) Representative confocal microscopy images of Pt1 and Pt9 cells expressing CrGES-mVenus in the diatom cytosol. The WT control is a non-transformed Pt1 strain. Scale bars represent 10 µm.

The differences we observed can arise from various factors, including algal biology, physiology and genetic variability, which may affect the compatibility of a certain strain with the conjugation protocol. For instance, in other techniques used for the transformation of diatoms, such as electroporation, the use of specific reagents like saponins, which facilitate the formation of membrane pores for DNA passage, significantly enhances the protocol’s effectiveness in algae where cholesterol is the main membrane component, such as *Nannochloropsis oceanica*. Additionally, synchronizing the life cycle has been shown to improve the efficiency of these protocols but while manipulating culture conditions (e.g., by changing light intensity) is known to improve transformation efficiency [84–86], in diatoms and other algae such as *Chlorella vulgaris* [87]. The conjugation protocol relies on long-term exposure to continuous light [6,35] which could favors Pt9 and Pt1 strains over Pt6 which would develop a stress response to light excess which would hinder the protocol success.

On the other hand, genetic variability among the three strains may help explain the observed differences in the success of transformation with our specific construct. Individual genome sequencing analyses determined that substantial genetic divergence exist among the ten classified *P. tricornutum* strains. Based on sequence similarity of the *ITS2* region within the 28S rDNA sequence, these strains can be divided into four clades. Clade A (Pt1, Pt2, Pt3, and Pt9), Clade B (Pt4), Clade C (Pt5 and Pt10), and Clade D (Pt6, Pt7, and Pt8), with Clades B and C being the most distant [16,88]. In the three strains investigated in this study, the zeocin resistance gene *sh-ble* was driven by the widely used promoter region of the gene *fucoxanthin chlorophyll a/ c binding protein B (FcpB, Phatr3_J25172)* and the emergence of zeocin-resistant colonies was dependent on the activity of this promoter. Conversely, the expression of the *CrGES-mVenus* fusion gene is driven by the promoter region of the *Phatr3_49202* gene, a recently identified strong constitutive regulatory sequence that is increasingly used for transgene expression in Pt1. [7,24]. The sequence of the *FcpB* promoter is nearly identical in the three strains, which could explain why all of them produced resistant colonies; however, *Phatr3_J49202p* shows a higher level of single nucleotide variation (SNV) among the strains (Fig. S6), potentially making the construct incompatible in some strains, as evidenced by the absence of mVenus signal in all Pt6 colonies.

Next, we set out to quantify the yield of heterologous geraniol in selected transgenic cell lines of the strains Pt1, Pt6, and Pt9, in 9 days batch cultivations both in CL and PP light regimes. Interestingly, geraniol yields were low (∼7.6 µg/L) and almost at the detection limit in the transformed Pt9 lines, regardless of the light regime. Conversely, the transformed Pt1 lines yielded higher amounts of geraniol with significant differences in the yield between cultures cultivated in CL (50.04 µg/L) and PP (34.21 µg/L) (Fig. 10c). The SNVs present in the promoter regions of *Phatr3_J49202* could have affected the expression of the *CrGES-mVenus* in Pt9, potentially resulting in low protein levels for significant geraniol production. This strain might also have lower cytosolic GPP levels compared to Pt1, which may not be adequate to support substantial geraniol production. The yields measured in Pt1 were lower than previously reported for the same strain transformed with the same constructs and presented differences dependent on light regimes, which were not observed before [26]. The overall lower yields could be due to slight differences in the cultivation equipment and parameters across different laboratories, and/or intrinsic metabolic differences resulting from strain adaptation and evolution to laboratory conditions in which strains are maintained. The differences observed between CL and PP in the yield of Pt1 transgenic cell lines could be due the longer cultivation time employed here (13 days) as compared to previous work, in which diatoms were cultivated for 10 days [26]. It should be noted that the heterologous monoterpenoid yields measured in these experiments are an underestimation of the actual production capacity. Geraniol is a volatile compound that easily escapes the culture medium and vessel. This experiment was not focused on the absolute and optimized capture and quantification of this compound, but rather on the relative biosynthetic capacity of the different strains. Hence, we grew the diatoms in an open shake system with an isopropyl myristate overlay. In such conditions, it is expected that a considerable fraction of the produced metabolite gets lost during the cultivation in open systems such as shake flasks, and during sampling. These observations underscore the importance of metabolic and genetic configurations in the success of metabolic engineering strategies [80]. Additionally, microalgae are known to release volatile organic compounds (VOCs) in response to various environmental stimuli, which may include stressful conditions or routine environmental factors such as light-dark cycles or seasonal changes [89]. Based on our previous observations, continuous light exposure may be more stressful for Pt1 than for Pt9, potentially inducing stress and increasing geraniol release as a response. This is similar to the behavior observed for isoprene and some monoterpenoids in other diatom species [90]. In this scenario, stressful conditions could have a beneficial impact on the production and release of this and related valuable compounds. This hypothesis is currently being investigated in our laboratory. We could also confirm that the transformed lines did not show a different growth or photosynthetic performance compared to the wild type strains cultivated in CL and PP (Fig 10 a,b). The growth pattern of the transgenic lines was similar, with higher cell density in CL compared to PP and higher growth for Pt9 over Pt1 in CL conditions. The photosynthetic activity followed the same trend observed for untransformed cells, confirming that PP is the optimal light regime for maintaining high photosynthetic yield in diatom cells (Fig 10 a,b) and at the same time confirming that the heterologous production of geraniol at these yields, not have an overloading effect on the overall diatom metabolism.

**Figure 10:**
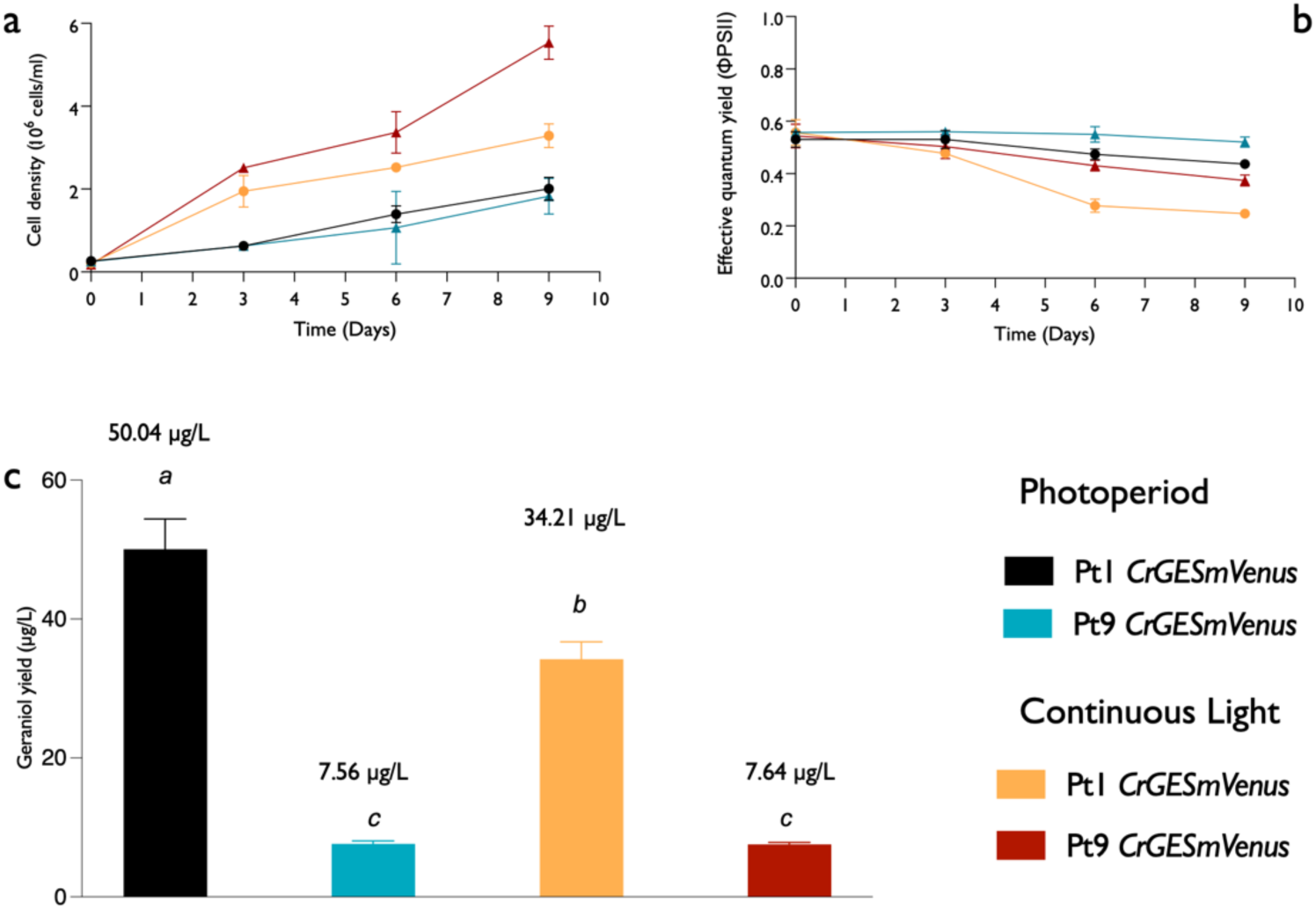
Profiling of transgenic Pt1 and Pt9 strains expressing *CrGES-mVenus* and cultivated either in PP or CL. Upper panels: effect of different light regimes on the (a) growth and (b) photosynthetic activity. (c) Production of heterologous geraniol in the transgenic strains grown in either PP or CL after 13 days of cultivation. Plots represent the estimated yields (µg/L) calculated based on the cell density at harvest. Values correspond to the mean of 3 independent replicates, error bars represent SD. Different letters indicate statistically significant differences between samples (one-way ANOVA test, *P*<0.001).

Finally, despite its relatively lower growth and photosynthetic efficiency, the Pt1 strain appears to be better suited for metabolic engineering and more efficient in producing heterologous monoterpenoids.

## Conclusions

This study provides a comprehensive evaluation of three ecologically distinct strains of *P. tricornutum* (Pt1, Pt6, and Pt9) under different light regimes to optimize their use as biotechnological platforms. The results highlight significant strain-specific differences in growth rates, photosynthetic efficiency, and pigment and triterpenoid amount and composition, emphasizing the critical role of light conditions in modulating these parameters. Pt9, originating from a tropical climate, exhibited superior growth and photosynthetic performance under continuous light, suggesting its potential for high-light environments. Pt6, adapted to intertidal zones, showed resilience under fluctuating light conditions, making it suitable for environments with variable light intensities. Moreover, the study identified Pt1 as the most effective strain for heterologous geraniol production, and, as such, the most suitable as a biotechnological platform. These findings underline the importance of selecting appropriate strains and light conditions tailored to their specific physiology for targeted metabolic engineering applications, thereby enhancing the production of valuable compounds. The insights gained from this research pave the way for more efficient exploitation of diatom strains in algae biotechnology, offering significant potential for commercial applications.

## Author contributions: CRediT

LM: Conceptualization, Investigation, Methodology, Data curation, Formal analysis, Writing – original draft, Writing – review and editing; PP: Investigation, Methodology, Formal analysis, Writing – review and editing; FP: Formal analysis, Writing – review and editing; MB: Investigation, Writing – review and editing; MF: Conceptualization, Methodology, Formal analysis, Writing – review and editing, Project administration, Resources, Supervision, Funding acquisition.

## Acknowledgements

This work was supported by the Novo Nordisk Foundation (grant number NNF22OC0078846 to MF), the Villum Fonden (grant number 37521 to MF) and an SDU Climate Cluster (SCC) with a Research Infrastructure Grant to MF. The authors would like to thank Lars Duelund for the technical assistance with analytical chemistry.

