## Supplementary Material File 1 for "Specific light-regime adaptations, terpenoid profiles and engineering potential in ecologically diverse *Phaeodactylum tricornutum* strains"

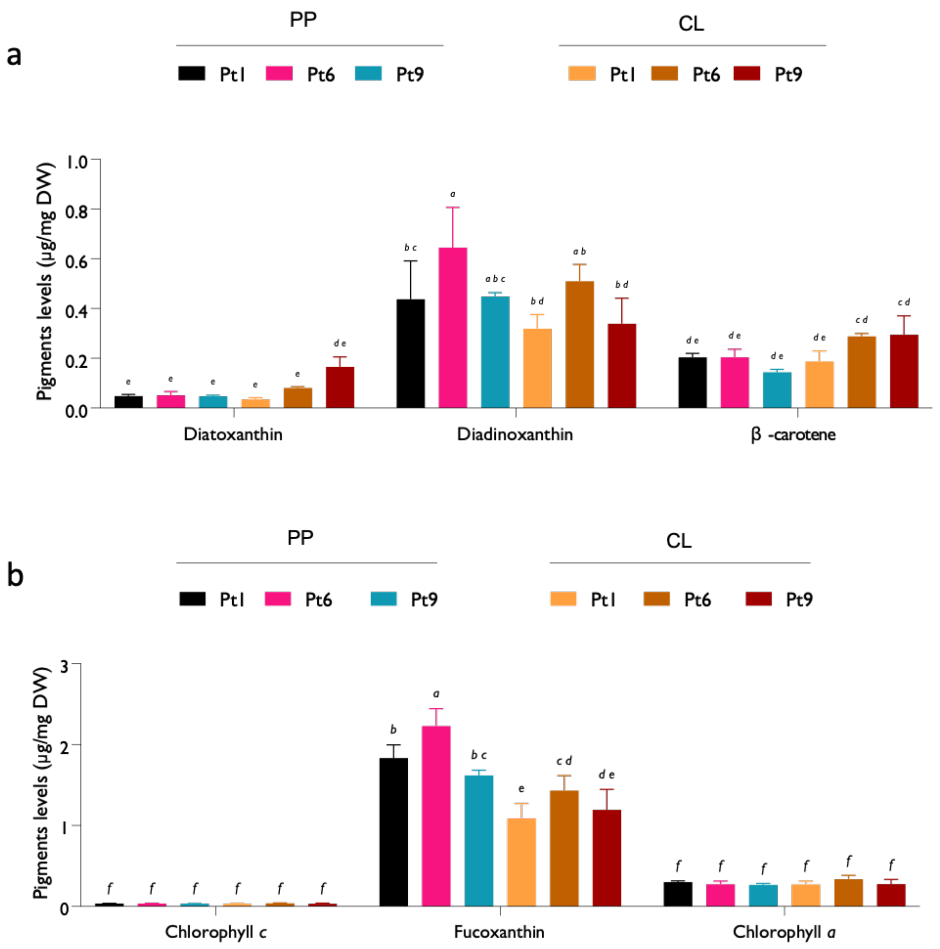


**Figure S1:** Levels of the single pigments in Pt1, Pt6 and Pt9 cultured under CL or PP. Pigments levels are expressed as µg/mg of dry weight and calculated according to a calibration curve of the authentic standards. Plots show the mean and SD of 3 independent replicates. Different letters indicate statistically significant differences between samples (one-way ANOVA, *P*<0.05).

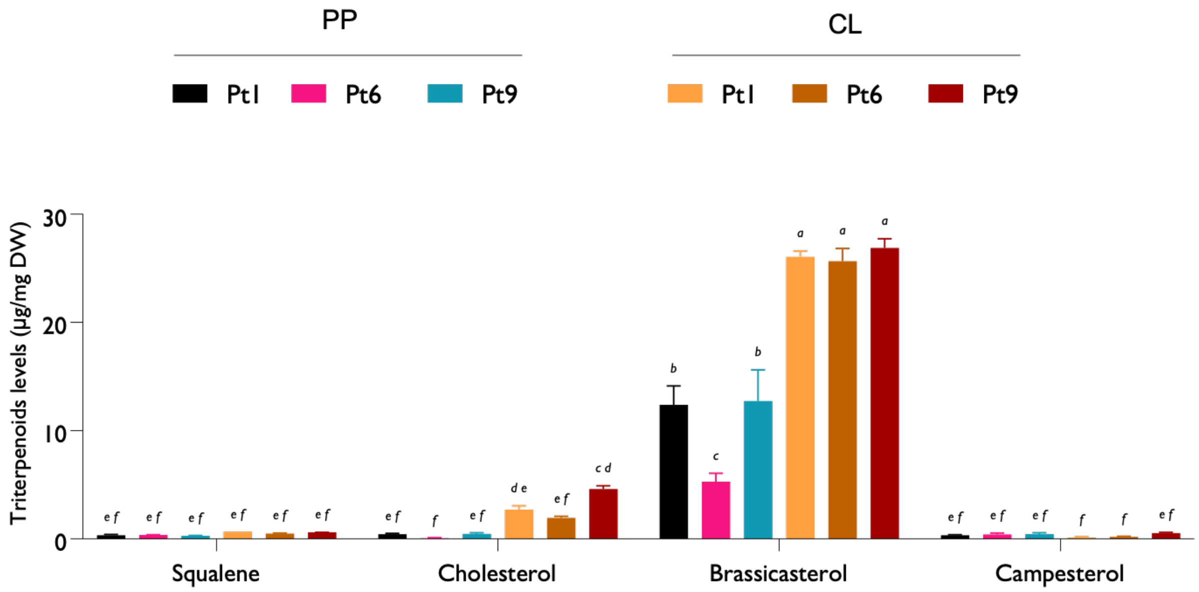


**Figure S2**: Levels of the single triterpenoids in Pt1, Pt6 and Pt9 cultured under CL or PP. The triterpenoid amount is expressed as µg/mg of dry weight and calculated according with a calibration curve of the authentic standards. Plots show the mean and SD of 3 independent replicates. Different letters indicate statistically significant differences between samples (one-way ANOVA, $P$<0.05).

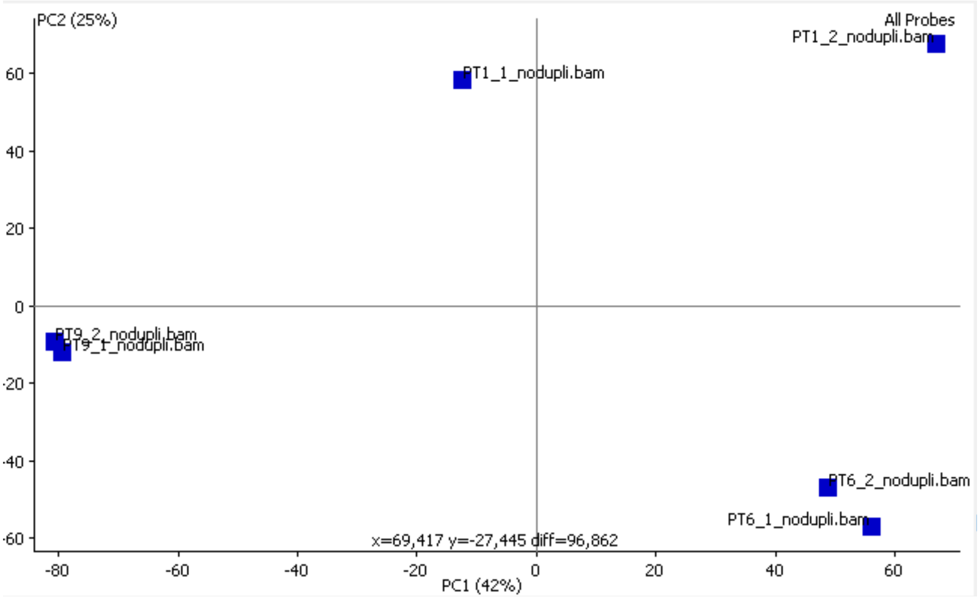


| | | R-squared |
|---|---|---|
| PT1_1 | PT1_2 | 0,756 |
| PT6_1 | PT6_2 | 0,8 |
| PT9_1 | PT9_2 | 0,99 |

**Figure S3:** Principal component analysis showing the variability among the 2 different replicates in the transcriptomic analysis by Chaumier et al. [34] and the corresponding R-squared values.

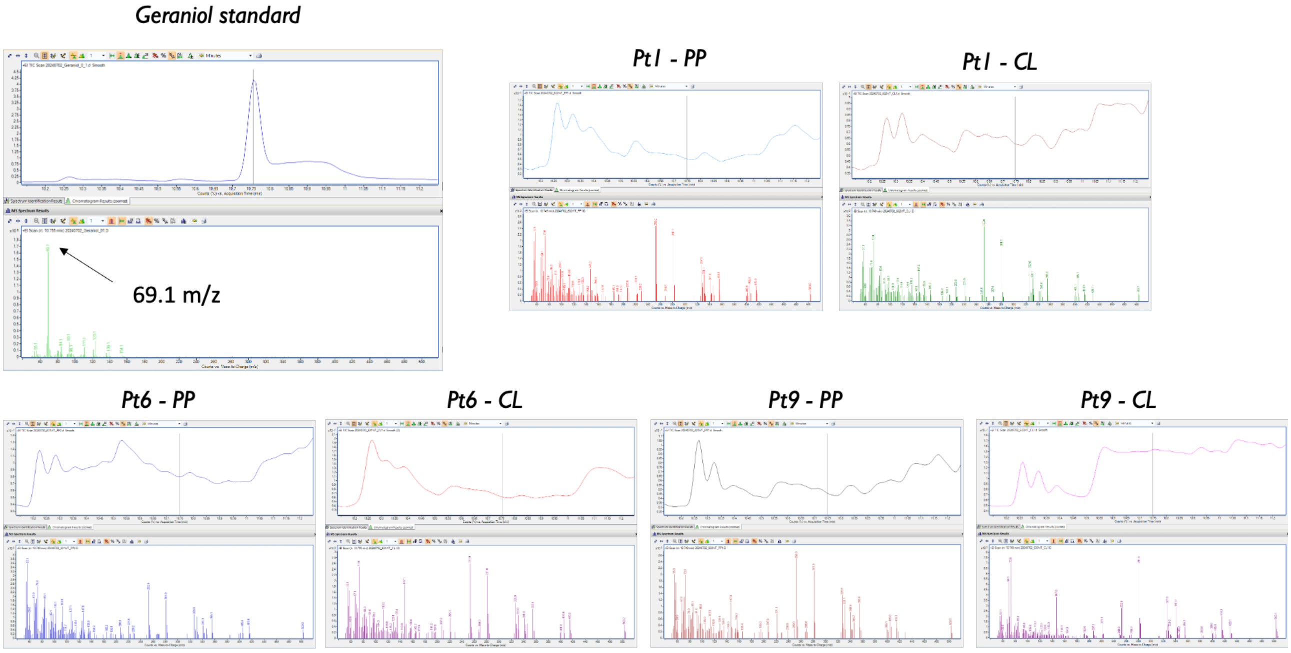


**Figure S4**: GC-MS chromatograms and mass spectra showing the fragmentation mass (69.1 m/z) of geraniol authentic standard and its absence in  wild-type Pt1, Pt6 and Pt9 strains.

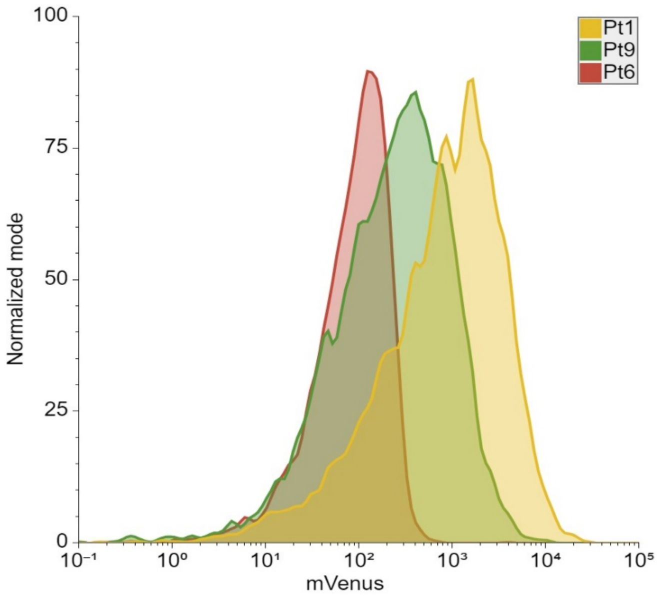


**Figure S5**: Representative histogram of Pt1, Pt6 and Pt9 with peak heights normalized to mode. Observations are representative 3 individual screenings each of them comprising the analysis of 5000 events. The plots were designed in Floreada.io.

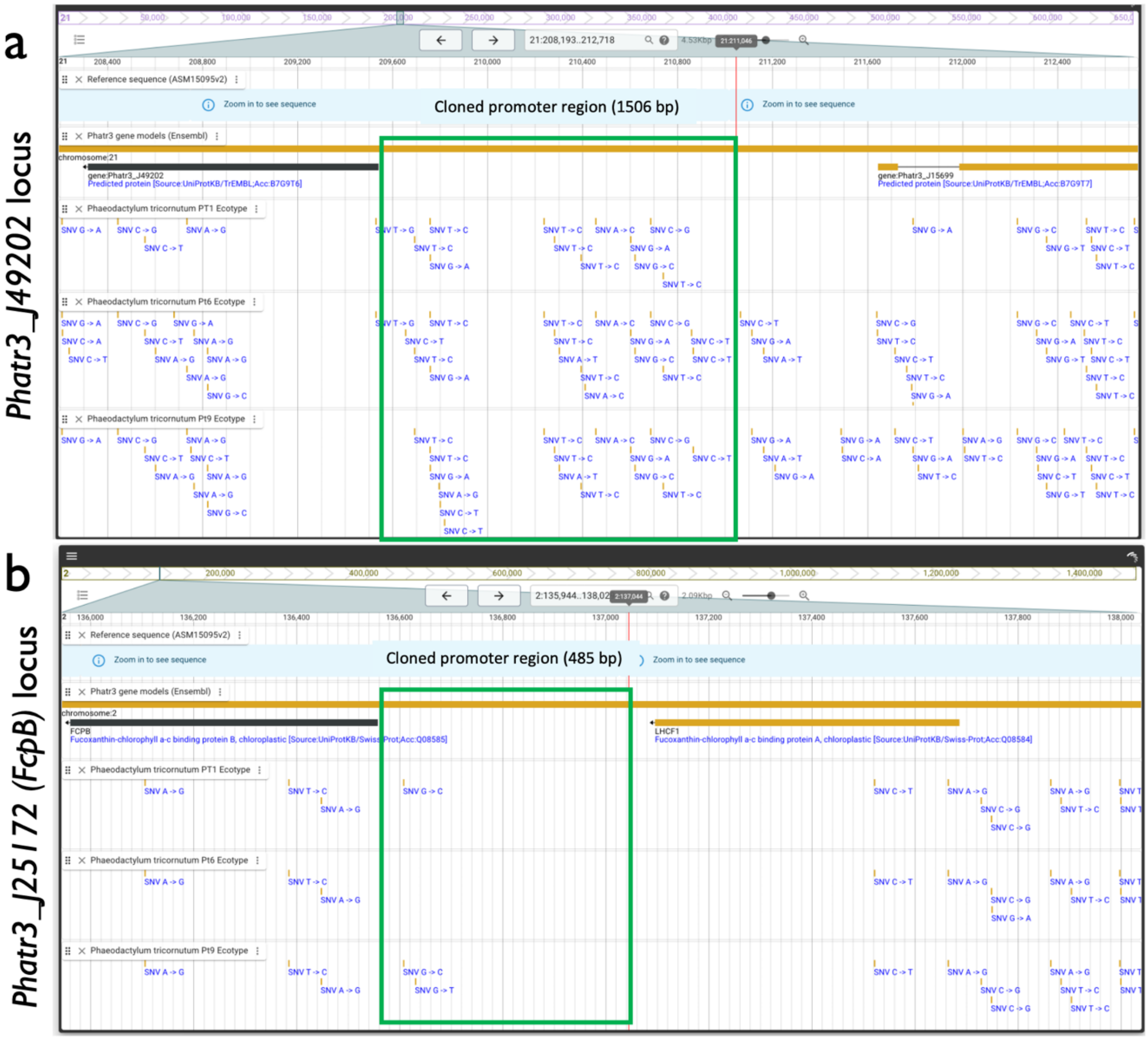


**Figure S6.** Overview of SNVs in the promoter regions (green boxes) of *Phatr3_J49202* (a) and *Phatr3_J25172* (*FcpB*) (b) of Pt1, Pt6, Pt9 compared to the reference genome assembly (ASM15095v2). Data obtained from DiatOmicsBase (https://www.diatomicsbase.bio.ens.psl.eu/).
